# Evaluation of abstract rule-based associations in the human premotor cortex during passive observation

**DOI:** 10.1101/2023.06.06.543581

**Authors:** Niloofar Gharesi, Lucie Luneau, John F. Kalaska, Sylvain Baillet

## Abstract

Decision-making often manifests in behavior, typically yielding overt motor actions. This complex process requires the registration of sensory information with one’s internal representation of the current context, before a categorical judgment of the most appropriate motor behavior can be issued. The construct concept of embodied decision-making encapsulates this sequence of complex processes, whereby behaviorally salient information from the environment is represented in an abstracted space of potential motor actions rather than only in an abstract cognitive “decision” space. Theoretical foundations and some empirical evidence account for support the involvement of premotor cortical circuits in embodied cognitive functions. Animal models show that premotor circuits participate in the registration and evaluation of actions performed by peers in social situations, that is, prior to controlling one’s voluntary movements guided by arbitrary stimulus-response rules. However, such evidence from human data is currently limited. Here we used time-resolved magnetoencephalography imaging to characterize activations of the premotor cortex as human participants observed arbitrary, non-biological visual stimuli that either respected or violated a simple stimulus-response association rule. The participants had learned this rule previously, either actively, by performing a motor task (active learning), or passively, by observing a computer perform the same task (passive learning). We discovered that the human premotor cortex is activated during the passive observation of the correct execution of a sequence of events according to a rule learned previously. Premotor activation also differs when the subjects observe incorrect stimulus sequences. These premotor effects are present even when the observed events are of a non-motor, abstract nature, and even when the stimulus-response association rule was learned via passive observations of a computer agent performing the task, without requiring overt motor actions from the human participant. We found evidence of these phenomena by tracking cortical beta-band signaling in temporal alignment with the observation of task events and behavior. We conclude that premotor cortical circuits that are typically engaged during voluntary motor behavior are also involved in the interpretation of events of a non-ecological, unfamiliar nature but related to a learned abstract rule. As such, the present study provides the first evidence of neurophysiological processes of embodied decision-making in human premotor circuits when the observed events do not involve motor actions of a third party.

## Introduction

We continuously receive and process sensory information to issue categorical judgments about how to best interact with our environment via motor actions to achieve certain goals (Gold and Shadlen, 2007; Cisek and Kalaska, 2010). In many contexts, we make covert categorical decisions that do not culminate in a companion motor action, such as when we want to assess whether an object is edible without tasting it. However, we may argue that *choosing not to act* is itself a sensorimotor decision (Muhammad et al., 2006; Wallis and Miller, 2003; Kalaska and Crammond, 1995). An alternative outcome might be to decide to perform an overt motor action, such as tasting that new unknown object. In that eventuality, the sensorimotor decision is transformed into a course of actions by which the motor system executes this plan to achieve the goal.

We may therefore conceive that sensorimotor decision-making proceeds through *sensory-perceptual, cognitive*, and *motor* processing stages in a specific behavioral context. These stages have long been regarded as functionally distinct, sequential, and implemented by functionally specialized brain circuits. More recently, however, a consensus has gradually emerged around the notion of *embodied* decision-making, whereby cortical motor circuits participate in both the *selection* and the *execution* of voluntary motor actions, even before a final action decision is made. Accordingly, behaviorally salient information is first extracted from sensory inputs and registered with internal representations of the context of the situation and expressed as evidence for actions in a space of motor representations of potential behaviors, before a decision of the action to be performed is made (Gold and Shadlen, 2007; Cisek and Kalaska, 2010; Cisek and Pastor-Bernier, 2014; Lepora and Pezzulo, 2015; Gordon et al., 2021). We sought to characterize in the present study the neurophysiological involvement of the human premotor cortex in embodied decision-making behavior.

There is extensive evidence that the premotor cortex (PM) is involved in the planning and execution of voluntary movements (Shenoy et al., 2013; Weinrich et al., 1984; Wise, 1985; Wise et al., 1997; Crammond and Kalaska, 2000). For example, single-unit neurophysiological recordings in the dorsal aspect of the premotor cortex (PMd) of non-human primates (NHPs) reflect the desired properties of an intended movement (e.g., target location, distance to target, movement direction and velocity) during an instructed delay period long before movement is initiated (Crammond and Kalaska, 1994; Weinrich et al., 1984; Mitz et al., 1991; Wise et al., 1997; Caminiti et al., 1991; Messier and Kalaska, 2000). Importantly, a wealth of NHP studies have shown that premotor circuits are also engaged in even earlier stages of motor planning, before the selection of a specific action, during the sensorimotor decision-making phase of tasks that involve arbitrary conditional stimulus-response association rules (Cisek and Kalaska, 2005; Thura and Cisek, 2014; Lepora and Pezzulo, 2015; Mitz et al., 1991; Muhammad et al., 2006; Rossi-Pool et al., 2016, 2017; Wallis and Miller, 2003; Wise et al., 1996, 1997; Hoshi and Tanji, 2002; Nakayama et al., 2008; Yamagata et al., 2009).

For instance, (Cisek and Kalaska, 2005) trained NHPs to choose between two reach targets based on a sequence of two visual cues and the application of a learned color-location conjunction rule. At the onset of a trial, two color-coded targets are displayed on opposite sides of a central starting arm position. This first visual cue informs the subject about the two potential actions that could be performed in that trial, but they need more information to determine which of the two targets will be its reach destination. During this first delay period, two separate populations of PMd neurons that are respectively associated with each target fire simultaneously, instantiating the concurrent representations of the two respective target-reaching actions. The onset of the second visual cue, a monochromatic circle at the central starting position whose color matched one of the two targets, determines the actual target in the current trial, but the subject is tasked to withhold its motor action until the onset of a subsequent “GO” cue. During that second delay period, although overt movements are not expressed yet, the firing of PMd neurons associated with the instructed target is enhanced while the other neural population is suppressed. These neurophysiological observations are compatible with PMd circuits instantiating, prior to movement onset, a representation of potential and final selected motor actions according to available sensory information and learned arbitrary stimulus-response association rules (Cisek and Kalaska, 2005, 2010).

Subsequently, (Coallier et al., 2015) used a multi-colored checkerboard instead of a monochromatic colored circle as the visual cue driving sensorimotor decisions. In each trial, the checkerboard comprised a different proportion of squares of the same two colors as those of the targets. That cue design controlled the level of conflicting evidence in favor of the decision to reach towards either of the targets. The NHP subjects were trained to decide in each trial which of the two colors was most prominent in the checkerboard cue, before reporting their perceptual decision via a reach movement towards the corresponding target. The data showed that the level of activity of PMd neurons was unaffected by the colors of the cues, but covaried strongly with the ambiguity of the evidence conveyed by the colored cue and by the duration of the time needed to make the color decision and initiate a reach movement to the chosen target. Finally, the PMd activity was strongly related to the direction of both correctly and incorrectly chosen targets, but not by their color. This study provided further support for a role of the premotor cortex in sensorimotor decision processes that rely on sensory evidence supporting the two target choices in the context of abstract target color-location conjunction rules, but makes little or no contribution to the perceptual decision about the dominant color of the checkerboard.

(Wang et al., 2019) used another task variant in which the checkerboard appeared before the two colored targets. The NHPs were tasked to make perceptual decisions about the dominant color of the checkerboard cue during the first delay period without being able to associate that decision with a specific reach movement (Coallier and Kalaska, 2014). Their results showed that PMd neural firing was weak during the first delay period, but again represented the direction of the decided movement only *after* the colored targets appeared to inform the subjects to which target they were to reach towards to report their decision about the checkerboard color cue. In an even more abstract fashion, other studies showed that NHP premotor circuits are also involved in the selection of reach targets based on their respective rewarding values and spatial separation (Pastor-Bernier and Cisek, 2011), including when subjects switch between targets of different values (Pastor-Bernier and Cisek, 2011), or when their target selection adjusts to fluctuating and probabilistic visual instructions (Thura and Cisek, 2014, 2016; Thura et al., 2022).

Taken together, these findings in non-human primates form a body of empirical evidence *i*) that premotor cortex activity reflects dynamic sensorimotor decision processes that are embodied in a *motor space* of possible movements, rather than in a *perceptual space* of abstract sensory evidence (here, based on color), and *ii*) that these processes of decision-making unfold *before* premotor circuits subserve the actual control of motor outcomes that stem from the decision (Cisek and Kalaska, 2010).

Once a chosen movement is performed by the subject, premotor and primary motor (M1) regions are also implicated in the *monitoring* of their movement execution errors. For instance, motor and premotor regions express single-unit and beta-band (15-35 Hz) activity whose magnitude reflects the size of reach errors during adaptation to perturbations by external force fields or by visuomotor rotations (Tan et al., 2014a,b; Torrecillos et al., 2015; Inoue et al., 2016). Motor and premotor beta-band activity typically decreases during the execution of motor actions, before expressing a transient *beta-rebounds* immediately after movement execution. The amplitude of beta rebounds is reduced when the motor plan is executed incorrectly, or when movements need to be readjusted in the current trial. This suggests that during motor skill learning, beta rebounds in premotor regions reflect the subject’s level of confidence about their internal predictive representations of motor plans during motor skill learning (Tan et al., 2016). Furthermore, a series of pioneering NHP studies demonstrated that premotor circuits are also involved in covert cognitive processes that do not lead to overt actions from the subject, including the recognition and understanding of the observed familiar or ecologically natural actions of a third party (another NHP or human agent) (Kemmerer, 2021; Cattaneo and Rizzolatti, 2009; Di Pellegrino et al., 1992; Gallese et al., 1996; Fogassi et al., 2005). They discovered that a sub-population of ventral premotor cortex (PMv, area F5) “mirror” neurons are activated both when a NHP executes a motor act such as grasping a pellet of food and when it observes a third party performing a similar action (Di Pellegrino et al., 1992; Gallese et al., 1996; Fogassi et al., 2005). A proposed interpretation is that the observer’s motor circuits instantiate a motor representation of the observed actions performed by the third party to facilitate their comprehension and evaluation (Kilner, 2011; Heyes, 2010; Kemmerer, 2021; Cattaneo and Rizzolatti, 2009; Rizzolatti et al., 1996). These neural representations may enable the observer to make causal interpretations of motor actions (Heyes, 2010), bridging across perception, cognition and action as an essential functional element for social interactions at large, and specifically for the assessment of the validity and the anticipation of the consequences of our own actions in a given context (Kilner, 2011; Heyes, 2010; Kemmerer, 2021; Cattaneo and Rizzolatti, 2009; Rizzolatti et al., 1996).

In humans, a large number of studies with electrophysiology, neurophysiology, and functional magnetic resonance imaging (fMRI) (Dinstein et al., 2007; Press et al., 2011, 2012; Rizzolatti et al., 1996; Grafton et al., 1996; Nishitani and Hari, 2000; Perani et al., 2001; Babiloni et al., 2002, 2016; Manthey et al., 2003; Tai et al., 2004; Muthukumaraswamy and Johnson, 2004; Koelewijn et al., 2008; Kilner et al., 2009a,b; Meyer et al., 2016; Del Vecchio et al., 2020; Fadiga et al., 1995; Hari et al., 1998; Järveläinen et al., 2001; Muthukumaraswamy et al., 2006; Cardellicchio et al., 2018) and behavioral psychophysics studies (Brass et al., 2001; Craighero et al.; Buccino et al., 2004; Paracampo et al., 2018; Avenanti et al., 2018; Press et al., 2005; Avenanti et al., 2018; Paracampo et al., 2018) have also shown that a broad set of cortical regions, including premotor and motor areas, respond to the observation of actions. The degree and nature of the observer’s personal experience with the observed movements, including the practice-dependent acquisition of new motor skills, are factors that modulate the degree of activation of their premotor system (Calvo-Merino et al., 2005; Orgs et al., 2008; Marshall et al., 2009; Cannon et al., 2014; Rüther et al., 2014; Meyer et al., 2016; Panasiti et al., 2016). These learning-dependent effects provide further support to the hypothesis that premotor activations during action observation reflect a form of embodied cognitive processing of observed events, through the covert neural simulation of observed actions from the individual motor repertoire of the observer (Kilner et al., 2004; Schubotz and Von Cramon, 2004; van Schie et al., 2004; Meyer et al., 2016; Avenanti et al., 2018; Paracampo et al., 2018).

Another body of studies has shown that premotor regions are also activated when participants evaluate the correctness of the observed outcomes of actions performed by a third party. It has long been known that electrophysiological markers of error detection such as the short-latency error-related negativity (ERN) and longer-latency error positivity (Pe) (Gehring et al., 1993; Dehaene et al., 1994; Carter et al., 1998), generated primarily in midline cingulate regions, both manifest when participants produce an incorrect motor output, or when they observe a human actor or avatar make such errors (van Schie et al., 2004; Panasiti et al., 2016; Pavone et al., 2016; Weller et al., 2018; Spinelli et al., 2018; Pezzetta et al., 2018; de Bruijn et al., 2007). However, more recent studies have shown that the passive observation of perturbed or incorrect actions of a third party also engages premotor and motor cortical circuits differentially, depending upon the correctness of observed actions (Manthey et al., 2003; Cardellicchio et al., 2018; Schubotz and Von Cramon, 2004; van Schie et al., 2004; Koelewijn et al., 2008; de Bruijn et al., 2007; Panasiti et al., 2016; Meyer et al., 2016; Malfait et al., 2010). For instance, (Manthey et al., 2003) reported fMRI effects in the premotor cortex of human participants watching an actor performing everyday activities either correctly or incorrectly (e.g., using an inappropriate tool). (Meyer et al., 2016) showed with EEG that the reduction of beta-band activity over the primary motor cortex is more pronounced when watching an actor clumsily grasping a target object. More abstract observed events also induce similar effects in premotor and motor cortex. For instance, (Koelewijn et al., 2008), using magnetoencephalography (MEG), reported decreased premotor and primary motor beta-band activity when participants are presented pictures of a human actor using abstract color-coded visual instructional cues to decide whether to press a right or left button with either their left or right finger. They showed that the temporal profile of this decrease of premotor and primary motor beta-band activity during task observation qualitatively replicated that of when the participants performed the task themselves, but was of significantly smaller amplitude. Importantly, beta-band rebound amplitude was larger when the participants made or observed incorrect finger or button presses, than when they made or observed correct responses.

The premotor cortex also responds when observing the actions of a diversity of non-human agents involved in motor tasks, such as digital avatars (Perani et al., 2001; Casile et al., 2010; Spinelli et al., 2018; Pezzetta et al., 2018), robotic and other automated agents (Tai et al., 2004; Costantini et al., 2005; Press et al., 2005; Gazzola et al., 2007; Oberman et al., 2007; Cannon et al., 2014; Rüther et al., 2014), and even abstract evocations of natural arm and hand movements with point-motion stimuli (Press et al., 2011; Ziccarelli et al., 2022; Kessler et al., 2006; Engel et al., 2008a,b).

These findings raise the question whether premotor cortex is also engaged in tasks in which participants observe non-biological stimuli that do not convey any inherent motor affordance and are not evocative of any obvious form of motor imagery. Some early studies indeed emphasized that premotor circuits do not respond to such arbitrary visual inputs on their own and that their activation requires that sensory information is related, either directly on via a learned association, to natural movements (Hari et al., 1998; Babiloni et al., 2002; Oberman et al., 2005; Costantini et al., 2005). For instance, (Cross et al., 2009a,b) showed that premotor circuits can be activated by arbitrary non-biological stimuli, but only after the observers are trained to associate these sensory events with specific outputs from their motor repertoire.

Other authors showed, however, that premotor activation does occur when processing arbitrary stimuli not previously associated with motor outputs, but only in the critical condition that the observers must make a categorical non-motor decision about the nature of the stimuli, and then report their decision by a motor act, like a simple button press. For instance, (Engel et al., 2008a) presented participants with a series of four short video clips per trial that consisted of either combination of arbitrary arm-waving gestures performed by a human actor or arbitrary animations of two spherical objects with approximately similar kinematics. The fMRI participants were asked to decide whether the fourth video was identical to one of the three preceding clips and report their decision with a button press. The videos featuring human movements presented during the first three clips activated a distributed set of cortical regions, including the bilateral ventral (PMv) and dorsal premotor (PMd) cortices, supplementary motor area (SMA), and the precentral gyrus. Those same regions were also activated, albeit at weaker levels, when any of the first three clips showed movements of spherical objects. The authors interpreted their data as showing that the arbitrary, yet human-like, motions of non-biological objects induced in the premotor circuits of the observers some cognitive associations with the equally arbitrary gestures of the human actor. In a companion study, the same authors used more abstract visual stimuli that consisted of three pseudo-randomly animated geometric shapes to minimize the possible confound of such cognitive associations with human movements (Engel et al., 2008b). In some of the trials, the participants were tasked to detect a color change of one of the symbols, and in others, to decide whether the motions could also be performed by human hands. Their hypothesis was that this latter categorical decision would require the participants to perform a covert simulation of their own hands moving to determine if they could perform the observed object motions. In both conditions, the participants reported their decisions with button presses. Both sets of videos activated several cortical sensorimotor regions, including PMv and PMd, with stronger amplitude effects when the decision required a covert simulation of hand movements.

In (Wang et al., 2015), EEG participants observed a Flanker-like task performed by “a partner in another room” (in reality, a computer). They were first presented with a 5-letter string whose middle letter determined the correct response, followed by a second visual stimulus that was a symbolic representation of the computer’s response choice. The participants were tasked to report orally whether the partner’s choice was correct or incorrect at the end of each trial. The data showed modulations of alpha-band activity (8-12 Hz) over M1, PMd and PMv during the task’s visual presentations, although they did not contain a visible actor or motor action.

Finally, von Cramon and colleagues performed a series of studies in which participants observed, in each trial, sequences of 8-12 non-biological visual stimuli that comprised arrays of icons of different shapes, colors and sizes, or a variety of color gradients (Schubotz and von Cramon, 2001, 2002a,b; Wolfensteller et al., 2007). The stimuli were designed to bear no inherent pragmatic value, did not suggest any form of transitive action, and did not offer any inherent motor affordance that would have evoked covert mental imagery of a corresponding action. The participants observed the sequences of visual stimuli, and identified their sequential structure based on different physical properties of the first several stimuli in the series, such as the physical identity of the icons, their spatial locations in the arrays, or the timing of their presentation. They were then tasked to detect and report with button presses in each trial whether one of the last stimuli in the series violated the correct sequential structure in the current trial. All these studies reported activation effects in several premotor and sensorimotor regions while the participants observed the initial stimuli in each sequence, prior to the appearance of the last critical images that would determine their decisions.

In summary, the literature contains conflicting reports of the presence or absence of premotor cortical activations when observing non-biological stimuli. The non-biological motion stimuli used by (Engel et al., 2008a,b) may have evoked a covert simulation of similar motions in premotor cortex, either by temporal association (Engel et al., 2008a) or by design (Engel et al., 2008b). Premotor activations occur during the visual presentation of unfamiliar or artificial motions when participants are tasked to make categorical non-motor decisions about certain physical properties of the stimulus or based on learned stimulus-response rules. In addition, in all those situations, including (Engel et al., 2008a,b), the participants were instructed to report their decisions via differential motor actions, either with oral responses or button presses. Therefore, such premotor activations during the decision-making sequence are likely to be driven by the visual input information in the context of a task requiring the application of arbitrary stimulus-response association rules to report the categorical non-motor decision via a sensorimotor output (c.f., (Mitz et al., 1991; Muhammad et al., 2006; Wallis and Miller, 2003; Hoshi and Tanji, 2002; Nakayama et al., 2008; Yamagata et al., 2009).

Furthermore, in most of those studies, the participants were tasked to make a categorical decision about whether they observed a discrepancy or error in the sequence of visual events. (Wang et al., 2015) found that when the computer takes the wrong decisions, EEG error-related markers (ERN and Pe) in human observers reflect that they interpreted the corresponding sequence of visual events correctly. However, premotor and motor alpha-band activity in the observer is modulated by the correctness of the performer’s decisions only when the task is performed by a human actor, not a computer (Manthey et al., 2003; Koelewijn et al., 2008; Meyer et al., 2016).

To the best of our knowledge, none of the studies with non-biological stimuli reported the human premotor activity while the participants observed the critical final stimuli in the sequence or activity associated with their categorical decisions about the sequential structures of the non-biological stimuli based on a previously-learned arbitrary rule. Therefore, whether premotor circuits only reflect the sensory evidence conveyed by the non-biological stimuli or also reflect the categorical decisions of the participants is still uncertain. A further unresolved question is whether the premotor cortex supports the application of stimulus-response rules that govern the outcome of observed non-biological events without any prior natural or learned association with specific movements, and without requiring overt limb movements to report the decisions.

A single-unit recording study in NHPs reported by (Cisek and Kalaska, 2004) is relevant to these questions. NHPs first learned to perform arm-reaching movements in response to color-coded visual cues according to a color-location conjunction rule (Cisek and Kalaska, 2005). After recording the responses of PMd neurons during the active arm-movement task, the NHPs were then put in a situation in which they observed the same visual cues on a computer monitor while an experimenter not visible to the subjects performed the task and moved the cursor randomly towards the correct or incorrect target across trials (Cisek and Kalaska, 2004). The NHPs could not perform the arm movements necessary to earn a reward and were not tasked to report whether the cursor moved towards the correct direction. Instead, they were rewarded at the end of the trials that were performed correctly by the experimenter. Therefore, the NHPs were motivated to observe the visual events on the monitor and were able to predict accurately whether they would receive a reward at the end of each trial (Cisek and Kalaska, 2004). During this passive-observation task, a large proportion of their PMd neurons replicated their discharge patterns evoked during the active reaching task (Cisek and Kalaska, 2005). Furthermore, consistent with the subjects’ predictive behavior, the PMd activity evoked by the visual instructional cues before the onset of cursor movements predicted the direction where the cursor should move according to the learned color-location conjunction rule. Therefore, this predictive activity in PMd was consistent with subsequent cursor movements only when it was moved towards the correct target. When the cursor was moved in the wrong direction, PMd neurons continued to signal the expected correct direction. These effects demonstrate that during the observation of task events, the NHP premotor cortex does not simply instantiate a passive sensory representation of stimulus events. Instead, they suggest that dorsal premotor circuits express a covert predictive process about the correct motor output to be expected from previously learned rules and the visual instructional cues of the current trial. These observations are consistent with a role of premotor circuits in the contextual prediction and interpretation of external events, and as such, are aligned with the broader theoretical constructs of active, perceptual inference and predictive coding (Sadaghiani et al., 2022; Lakatos et al., 2009; Ballard et al., 1997).

Consistent with this perspective, (Cisek and Kalaska, 2004) proposed that premotor activity during the observation of instructional cues reflect the covert mental rehearsal or simulation of the movements that the subjects had learned to perform to move the cursor to the target specified by the visual cues (see also (Engel et al., 2008b,a; Cross et al., 2009a,b)). Through such internal representations, the subjects would be able to predict whether they will receive the reward if the cursor moves in the correct direction. Consistent with that assumption, the predictive activity in PMd ceased a few trials after the reward delivery system was inactivated and resumed a few trials after reward was re-activated (Cisek and Kalaska, 2004).

These findings provide further evidence that embodied sensorimotor decision-making processes in PMd are expressed in response to arbitrary visual cues and stimulus-response rules (Cisek and Kalaska, 2004, 2005, 2010; Wang et al., 2019). Those processes are engaged both when NHP subjects actively produce movements to obtain a reward, and when they passively observe another agent produce such movements and expect a free reward from the correct execution of the task rules. Therefore, premotor circuits are not only implicated in the application of learned arbitrary stimulus-response association rules to select and control overt voluntary arm movements, but are also recruited in covert predictive non-motor cognitive processes to assess the potential outcomes of observed external events.

Two essential open questions remain: 1) are premotor circuits also involved when subjects learn the arbitrary stimulus-response association rules under circumstances where the external events are not related to specific *motor* outputs? 2) are these phenomena expressed by the human brain? The present study reports on the involvement of human dorsal premotor activity when participants either i) perform a hand-movement task to displace a cursor towards one of two opposite targets, based on color-coded instructional cues interpreted via a simple color-location conjunction rule learned previously (Cisek and Kalaska, 2005) (Active Performance condition), or ii) passively observe the same visual events while a computer agent performs the same task with correct and incorrect outcomes (Cisek and Kalaska, 2004) (Passive Observation condition). We used non-invasive magnetoencephalography (MEG) cortical imaging Baillet (2017) as participants actively performed the task or passively observed the task being performed by a computer. The participants were split into two groups depending on how they learned the color-location conjunction rule that determined correct active or passive task performance. The *Active Learners* group actively performed the hand-movement task themselves and learned its rules by trial and error from feedback received after each learning trial. The *Passive Learners* group watched a computer perform the task correctly or incorrectly, as indicated via visual feedback on each trial and therefore learned the task rules without any association to specific hand movements. We expected to observe beta-band premotor activity unfold across the successive epochs of a trial, from the presentation of the color cues, the selection of the correct target, the preparation for and execution of hand movements or the evaluation of the computer’s decisions (Kilavik et al., 2013; Koelewijn et al., 2008). Specifically, we anticipated premotor activity when, following instructional cues, the participants make the correct decision about the overt hand movements they are about to perform in the Active Performance condition (Kilavik et al., 2013). We also predicted differential premotor effects in the Passive Observation condition, depending on the correctness of the computer’s decision (Koelewijn et al., 2008; van Schie et al., 2004). These hypotheses stem from the construct discussed above that the human premotor cortex is engaged in the appraisal of sensory information, even when of an arbitrary and non-biological nature. We also aimed to address the critical question whether during passive observation, the human premotor cortex is activated before or after the participants have learned to associate the visual stimuli with specific hand movements. If before, this would demonstrate that the human premotor cortex is also involved in cognitive decision-making. If after, the data would then show that the human premotor cortex is engaged in a more specific form of embodied decision-making linked to the observer’s acquired motor repertoire. More broadly, if there is no significant premotor effect during the passive observation of task events in both groups, this would suggest that the human premotor cortex bears a role in cognitive decision-making that is restricted to when observing ecological and familiar movements performed by a human agent.

## Materials and Methods

### Participants

A total of 18 healthy individuals (9 men, 9 women, age 27.2 *±* 4 years) with no prior history of neurological and psychiatric disorders participated in the study. All participants were right-handed and had normal or corrected-to-normal vision. Experimental protocols were approved by the Institutional Review Boards of the Montreal Neurological Institute and the Université of de Montréal Health Research Ethics Committee (CERES - Comité d’éthique de la recherche en santé de l’Université de Montréal) and respected all institutional and national guidelines. All participants provided written informed consent prior to their participation. All task conditions were performed inside the magnetically-shielded room of the MEG unit of McConnell Brain Imaging Centre, located at the Montreal Neurological Institute.

### Active performance of the two-target forced-alternative choice task

Visual stimuli were back-projected on a semi-transparent screen placed 50-70 cm away from the participants. In the Active Performance condition, the participants performed a two-target (2T) task (Cisek and Kalaska, 2004, 2005), which is a two-alternative forced-choice task where a color-coded target location is designated from eight possible locations, following two successive instructional cues (Fig. 1). A trial started with the display of a white circle (1.5cm diameter) at the center of the screen. The participants used a joystick with their right hand to position the on-screen cursor within that circle during a central fixation period of 500ms (center-hold-time [*CHT*] epoch). A green and a red circle (2 cm radius) were then displayed at two opposite target locations (8cm apart) of the eight possible locations during 1000ms (spatial-cue [*SC*] epoch). The participants were instructed to keep the cursor still within the central white circle during this first instructed-delay period. Next, the color-coded spatial cues were removed, requiring the participants to remember the color-location conjunction combination of the two spatial cues for 500ms (spatial-cue-off [*SCO*] epoch). The color of the central circle then turned to either red or green, designating which of the two memorized color-coded spatial cue locations is the correct target in the current trial (color-cue [*CC*] epoch). The color-cue instructed-delay period lasted 1000-1500ms, during which the participants continued to hold the cursor still inside the central circle. Next, eight white circles appeared around the central circle at each of the eight possible target locations (GO cue). The participants then used the joystick to move a cursor arrow outside the central circle towards the target location they had chosen. The trial counted as correct when the participants positioned the tip of the cursor arrow within the circle at the target location specified by the combination of the current spatial and color cues within 1500ms (movement-time [*MVT*] epoch) and held the cursor inside that target circle for another 3 to 4 seconds (target-hold-time [*THT*] epoch). All possible combinations of target color-location conjunctions were presented across trials in a balanced pseudo-random task design. To minimize eye movements, the participants were instructed to fixate their eyes on a crosshair at the center of the screen throughout each trial.

**Fig. 1.**
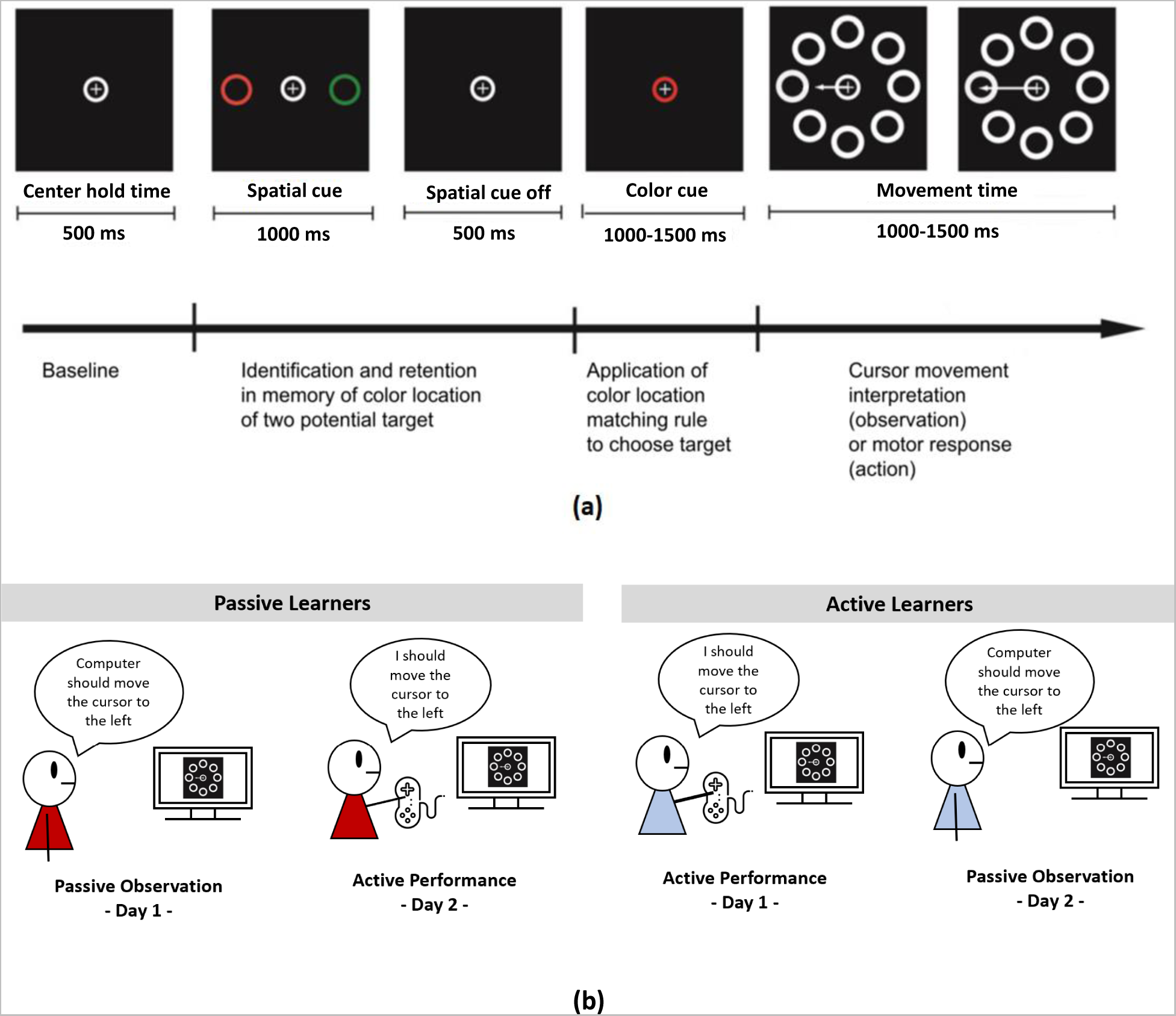
(a) Sequence of visual stimulus events in the 2T task, for a trial in which the correct target is to the left of the central starting position. (b) Cartoon representation of the two groups: Active Learners and Passive Learners, and tasks: Active Performance and Passive Observation.

### Passive observation of the two-target forced-alternative choice task

In the Passive Observation condition, the participants were presented the same sequences of visual stimulus events as in the Active Performance condition. However, they were instructed to keep their arms still and relaxed at their sides and did not hold onto or use the joystick. Instead, the presentation computer controlled the movements of the cursor. Following the successive presentation of the spatial and colors cues, at the onset of the GO cue, the computer extended the arrow out from the central window. In 50% of the trials, the arrow extended towards the correct target location, while in the other half of the trials, it extended the arrow towards the incorrect, opposite target location. All possible combinations of target color-location conjunctions, correct colored-target choices, and correct versus incorrect target choices were presented across trials, in a balanced pseudo-random task design. Here too, the participants were asked to fixate their eyes on a central crosshair.

### Passive vs. active learning tasks

The participants assigned to the Passive Learners group learned the task rules by watching the computer perform the task in the Passive Observation condition. Active Learners learned the task by trial-and-error while performing the task using the joystick in the Active Performance condition.

### Study design

All participants performed the 2T task in the Active Performance and Passive Observation conditions over two separate and non-consecutive visits. Passive Learners performed the task in the Passive Observation condition at their first visit (Fig. 1b). At their second visit, they performed the task in the Active Performance condition.

Reciprocally, Active Learners first performed the task in the Active Performance condition (Fig. 1b), and then in the Passive Observation condition at their second visit.

Both visits comprised three main phases including “task rule learning”, “task verification/familiarization” and “full data collection” (Fig. 2). MEG data were collected during both visits in both groups of participants.

**Fig. 2.**
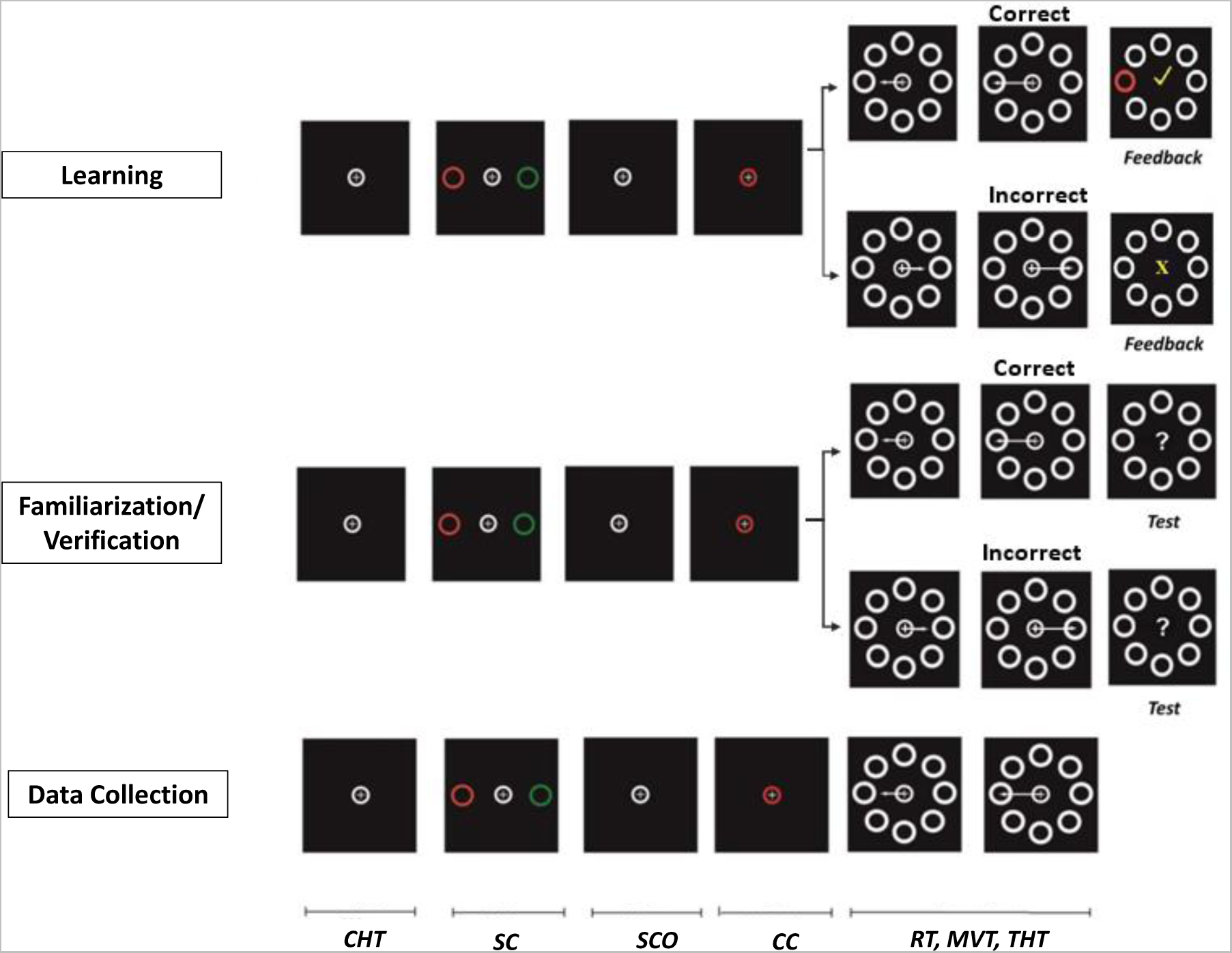
Sequences of stimulus events used for the learning, verification/familiarization, and data collection. During the familiarization/verification phase of the Passive Observation condition, the experimenter asked the participants to count the number of either correct or incorrect target choices made by the computer when they saw a question mark displayed after the last trial; they reported their count at the end of the 32-trial block. In the Active Performance condition, the question mark prompted the participants to count whether they had selected either the red or green target; they reported their count at the end of the 32-trial block. The reaction time (RT) is the time to movement onset after the display of the GO cue. The duration of the movement (*MVT*) is the time between movement onset and end. The target-hold-time (*THT*), is the time required to hold the cursor within the target (3 to 4 s).

#### Task rule learning and verification/familiarization

At the beginning of the first visit, the participants learned the 2T task either by Active Performance or Passive Observation. One sequence of stimulus events during a single trial of the learning phase is shown in Fig. 2, top row. One sequence of stimulus events during a single trial of the verification/familiarization phase is shown in Fig. 2, second row.

##### Passive Learners, Passive Observation

Passive Learners learned the color/location conjunction rule with visual feedback indicating whether the arrow controlled by the computer pointed toward the correct or incorrect target location in each Passive Observation trial. The feedback consisted of a white checkmark [✓] after correct trials and a yellow [X] mark after incorrect trials. Each block consisted of 32 trials. After each block, the participants were asked to share with the investigator what they thought they understood about the task rule. If they failed to explain the color-location conjunction rule adequately, they were presented with a second rule-learning block. Most participants were able to identify the rule within a single 32-trial block, and proceeded with the verification/familiarization phase (Fig. 2, second row), where they were asked to count the number of either correct or incorrect trials performed by the computer without receiving knowledge-of-results visual feedback at the end of each trial. If they succeeded in completing this phase satisfactorily, they moved on to the full data collection phase (see below).

##### Active Learners, Active Performance

Active Learners learned the color/location conjunction rule that determined correct outcomes by active trial-and-error performance of the 2T task using the hand-controlled joystick. After they placed the cursor in the center target, the sequence of *CHT, SC, SCO* and *CC* epochs unfolded, followed by the display of a GO cue. The participants then used the joystick to move the cursor to their chosen target location and keep it there for 1500ms. They received the same knowledge-of-results feedback than Passive Learners at the end of each trial. To engage their vigilance, the participants were also tasked to report at the end of the trial block how many times they had selected either the red or green target. As with Passive Learners, they were asked to explain their understanding of the 2T task rule after the first training block. They were offered a second block of trials, if necessary. Once they completed the verification/familiarization block with a minimum success rate of 75%, the participants proceeded to the full data collection phase. After the learning phase, all participants were able to complete the task in the Active Performance condition with 100% accuracy.

#### Full data collection

Following the verification/familiarization phase, the participants performed the task without further knowledge-of-results feedback. In the Passive Observation condition, they were instructed to count either the number of observed correct or incorrect trials in different trial blocks and to report their count at the end of each block (van Schie et al., 2004; Koelewijn et al., 2008). The counting instruction varied randomly between trial blocks. In the Active Performance condition, the participants counted and reported how many times they selected either the red or green target at the end of each trial block. The participants performed 4 blocks of 75 trials in the Passive Observation condition, and 4 blocks of 50 trials in the Active Performance condition.

Counting correct or incorrect trials in the Passive Observation condition required the participants to assess whether the computer did apply the color-conjunction rule correctly in each trial and to make a categorical correct/incorrect decision. Counting red or green targets in the Active Performance condition required the participants to categorize and count individual trials according to their color choice. Counting also assured that all participants remained attentively engaged in the task throughout data collection.

During the full data collection phase of the first visit, the Passive (respectively Active) Learners performed the task in the Passive Observation (respectively, Active Performance) condition (Fig. 1b). During the second visit, each group performed the task in the other condition from their first visit (Fig. 1b). Passive Learners applied the passively-learned task rule in the Active Performance conditions; Active Learners observed the computer’s performance in the Passive Observation condition. Both groups completed the same learning and verification/familiarization phases on their first visit, before going through four trial blocks in either the Passive Observation or Active Performance condition.

### MEG data acquisition

The participants underwent a T1-weighted structural MRI scan for subsequent MEG source mapping analyses. The MRI volume was realigned with the MEG data using three fiducial points (nasion and the left and right pre-auricular points) digitized at each MEG visit using a Polhemus 3-D digitizer (Polhemus Isotrak, Polhemus Inc., VT, USA). The alignment was refined using 100 additional digitized scalp points with Brainstorm (Tadel et al., 2011) .

We used FreeSurfer to obtain surface tessellations of the scalp and cortical surfaces from the individual MR volumes, with default parameter settings, which we then imported in Brainstorm and down-sampled to 15,000 cortical vertices for subsequent MEG source mapping (Baillet et al., 2001).

MEG data were collected with participants seating under a 275-channel VSM/CTF MEG system (anti-aliasing low-pass filter at 600Hz; sampling rate, 2400Hz) in a magnetically-shielded room (3-layer passive shielding). Electro-oculogram (EOG) electrodes were placed at the outer corners of the two eyes, and above and below the left eye to record horizontal (HEOG) and vertical (VEOG) eye movements, respectively, to detect eye blink and large saccades. Similarly, to detect and remove MEG artifacts produced by heartbeats, the electrocardiogram (ECG) was measured as a reference signal with a pair of electrodes across the chest below the clavicles with the same sampling frequency as MEG.

### MEG data preprocessing

All MEG data preprocessing and source imaging was performed with Brainstorm (Tadel et al., 2011); using default parameters) and custom-written scripts in Matlab (MathWorks, Natick, MA, USA). All implementation details are readily documented and can be verified in Brainstorm’s code. The MEG data were collected with built-in CTF’s 3rd-order gradient compensation. Jittered delays between the task presentation computer and monitor visual displays were registered for each trial by a photodiode located on the monitor in the MEG room. Power line noise (60 Hz and harmonics) was attenuated with 3-dB bandwidth notch filters. Sensors with poor quality were labeled as bad channels after visually inspecting the power spectrum of the signals. To clean the data from heartbeats, eye blinks, saccade contamination, and muscle artifacts, we designed specific signal-space projectors (SSP), following good-practice guidelines (Gross et al., 2013). The contaminated data segments were marked automatically based on signal variance observed in specific frequency ranges (e.g., 1.5-15 Hz and 10-40 Hz for eye blinks and heartbeats, respectively). The SSPs were derived from predefined time windows (e.g., 400-ms time windows centered around eye blinks, or 160-ms around each heartbeat event). Finally, the recordings were manually screened to identify any additional segments contaminated by head movements or environmental noise sources, which were excluded from subsequent analysis.

### MEG source imaging

We obtained a forward head model for each participant using the overlapping-spheres approach. We then derived weighted minimum-norm estimators of cortical source time series with constrained dipole orientations, all using Brainstorm with default parameters. We produced a noise covariance matrix from MEG empty-room recordings collected at each visit to account for environmental and system noise in the source mapping procedure. We defined a set of dorsal premotor cortex regions of interest (ROIs) in each hemisphere of the individual MRI, as the junction of the precentral and the superior frontal sulci, about 1.5cm anterior to the primary motor cortex, and the inverted omega shaped “hand knobs” of the dorsal part of the central sulcus (Fig. 8c) (Picard and Strick, 2001; Donner et al., 2009; Yazici et al., 2019).

### Time-resolved multivariate decoding

Prior studies have reported the decoding of movement direction and velocity from MEG sensor signals in performing wrist movements (Waldert et al., 2008; Wang et al., 2010; Jerbi et al., 2011, 2007). We replicated and extended this approach to decode the intended direction of the movement in its preparatory phase, prior to movement onset and in the absence of overt motor activity.

We derived MEG signal decoders to determine the movement direction i) during the preparation and performance of hand movements in the Active Performance condition, and ii) after the presentation of the information that specified the target location and during the Passive Observation of arrow movements. We epoched the MEG data in segments around each task cue, namely *SC, SCO, CC* and GO. We hypothesized that the color cue (*CC*) epoch is when the participants have accumulated complete evidence to determine their target in the current trial in both the Active Performance and Passive Observation conditions, and after which they prepared to execute a specific motor output plan in the Active Performance condition (Cisek and Kalaska, 2005). Therefore, we split the trials based on the intended (Active Performance) or observed (Passive Observation) movement direction.

From the 8 possible directions of the cursor movements (right, left, up, down, and the four diagonals), we pooled all trials in two categories whether the intended or observed arrow motion was directed towards the right of left hemifield of the display (Condition R and Condition L, respectively; Fig. 3). We discarded the trials where the targets were set at the Up and Down locations.

**Fig. 3.**
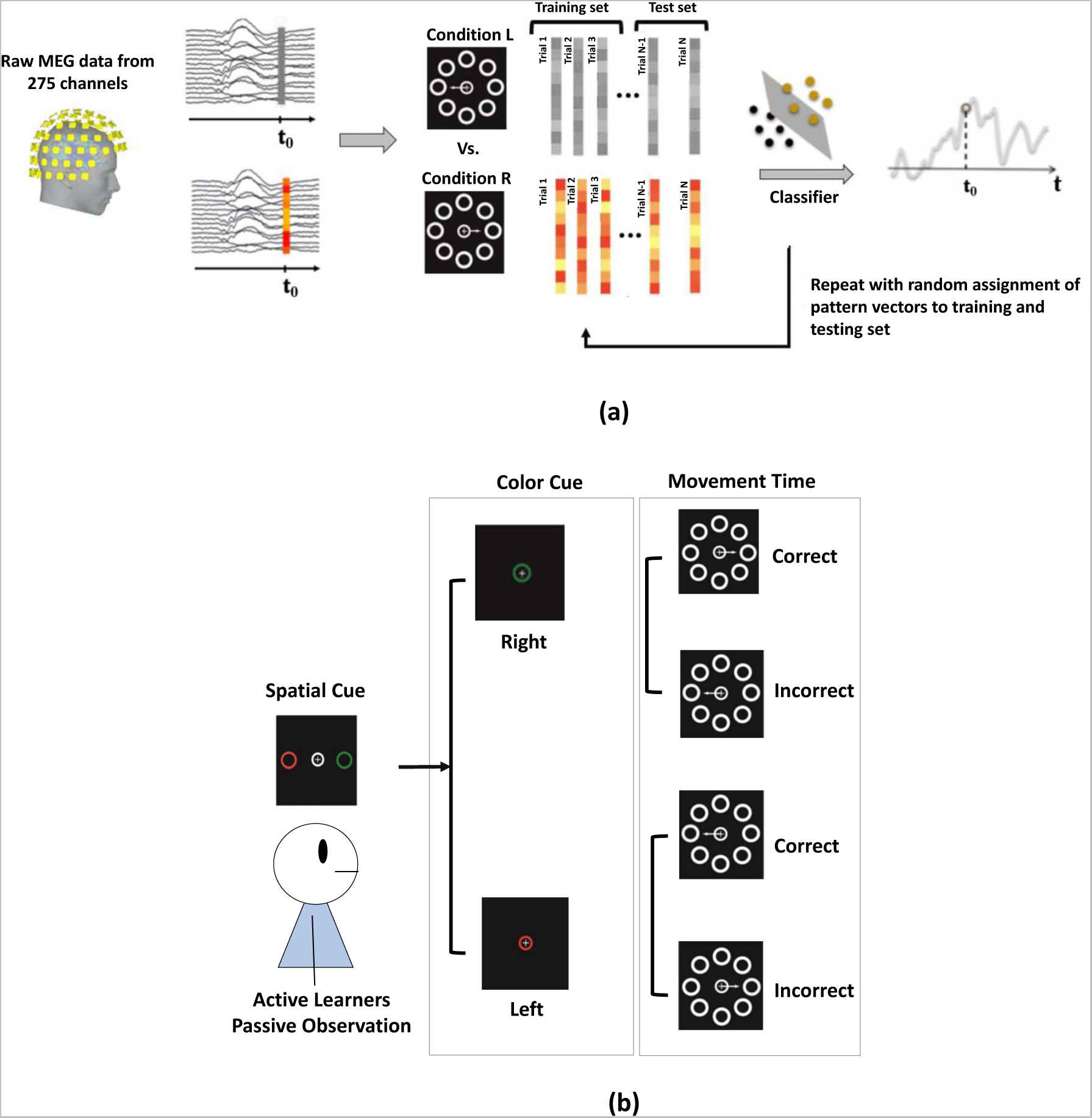
Multivariate decoding of MEG sensor data: (a) Time-resolved multivariate decoding was conducted using pairwise SVM classification at each time step. (b) Data clusters for the two trial Conditions (L, R) in the Passive Observation condition. The participants were tasked to observe the movements of the cursor as controlled by a computer. Their expectations of the cursor motion direction during the CC epoch could be similar or different from what they actually observed during the subsequent *MVT* epoch.

Because both the Passive and Active Learners moved the cursor to their target of choice in the Active Performance condition, the participants’ expectation (before movement onset) and observation of the direction of the actual arrow movement were always consistent. The participants’ decisions of the target location were always correct. However, in the Passive Observation condition, the target location decided by the computer was independent of the participants’ appraisal of the correct target location. Therefore, the computer-controlled arrow could be expected to move either towards the correct (as expected by the participants) or the incorrect target location.

We trained multivariate decoders of MEG brain signals to identify if and when i) the representation of the expected (Passive Observation) or intended (Active Performance) direction of movements towards the selected target location is expressed during the *CC* epoch, ii) the brain representation of the direction of the observed (Passive Observation) or executed (Active Performance) arrow movements is expressed during the *MVT* epoch, and iii) the representation of the correctness of the arrow movement direction is expressed, also during the *MVT* epoch, in the Passive Observation condition.

We used custom, Brainstorm and LIBSVM (cha) library functions for implementing the decoders based on linear support vector machines (SVM). The classification was conducted for each participant separately in a time-resolved manner, with independent decoders at each time point (Fig. 3). At each time step *t* of a trial in the given epoch of the task (Fig. 3a, left), the data consisted of a 275*×*1 vector of MEG sensor signals. We obtained such data vectors for all time steps and all trials of both the Condition Right and Condition Left target locations, yielding two arrays of 275*×N* vectors for each time step of the CC epoch: one set for intended or expected Left movements (Fig. 3a, center; grey columns) and one set for intended or expected Right movements (Fig. 3a center; colored columns), where *N* is the number of trials in each condition.

A subset of the data in Conditions Left and Right were selected randomly to train the SVM classifier and define a hyperplane that optimally separated the data vectors from the two task conditions (Figure 5a center; black and yellow dots: the 275*×*1 vectors for the training trials for Conditions Left and Right, respectively). The classifier was then tested on the remaining trials not used from training. We defined the decoding accuracy (DA) at each time step as the ratio of test trials correctly assigned to Condition Left or Right. For each participant, we obtained a DA time series using 10-fold cross-validation, where the trials used for training and testing the decoders were mutually exclusive and randomly bootstrap-sampled for each cross-validation iteration (we used 10 iterations ;Fig. 3a, right). The individual DA time series were then averaged across participants. The same classification procedure was used with different combinations of trials across the *CC* and *MVT* epochs (Fig. 3b) in the Active Performance and Passive Observation conditions.

### Extraction of beta-band activity

Using the FieldTrip plugins of Brainstorm, we applied multi-tapers(Mitra and Pesaran, 1999) over overlapping sliding time window of 1-s duration in steps of 100ms to obtain spectrograms of the cortical source time series in the 5-40 Hz frequency range (1-Hz step). The time windows were tapered with a Hanning window with a modulation factor of 10. From these spectrograms, we extracted the time fluctuations of cortical activity in the beta frequency band (15-29 Hz), including in the premotor regions of interest (PMd).

Fig. 8d shows the timeline of premotor beta-band activity in Active and Passive Learners. We standardized these time fluctuations using the z-score transform with respect to several baseline data segments:

1) 500 to 200ms prior to the *CHT* cue (Fig. 8d) ; 2) 500 to 200ms prior to the onset of the spatial cue (*SC*) (Fig. 9) ;

3) 1000ms to 200ms prior to movement onset (Figs. 10).

## Results

### Behavior

We measured the duration of the movement of the arrow (from onset to offset) in each trial and participant (Fig. 4). We found a small but significant difference between the mean duration of movements of Passive Learners (1.12 *±* 0.99 s) vs. Active Learners (1.36 *±* 1.15 s) in the Active Performance condition (paired two-sample t-test, *p <* 0.001). In the Passive Observation condition, because the movements were executed by a computer following a pseudo-random distribution, there was no significant difference in movement duration between groups. In that condition, half of the trials completed by the computer were incorrect (128 incorrect trials per participant on average). The participants always choose the correct target location in the Active Performance condition. This confirmed that they knew the color-location conjunction rule very well and were highly vigilant while performing the task.

**Fig. 4.**
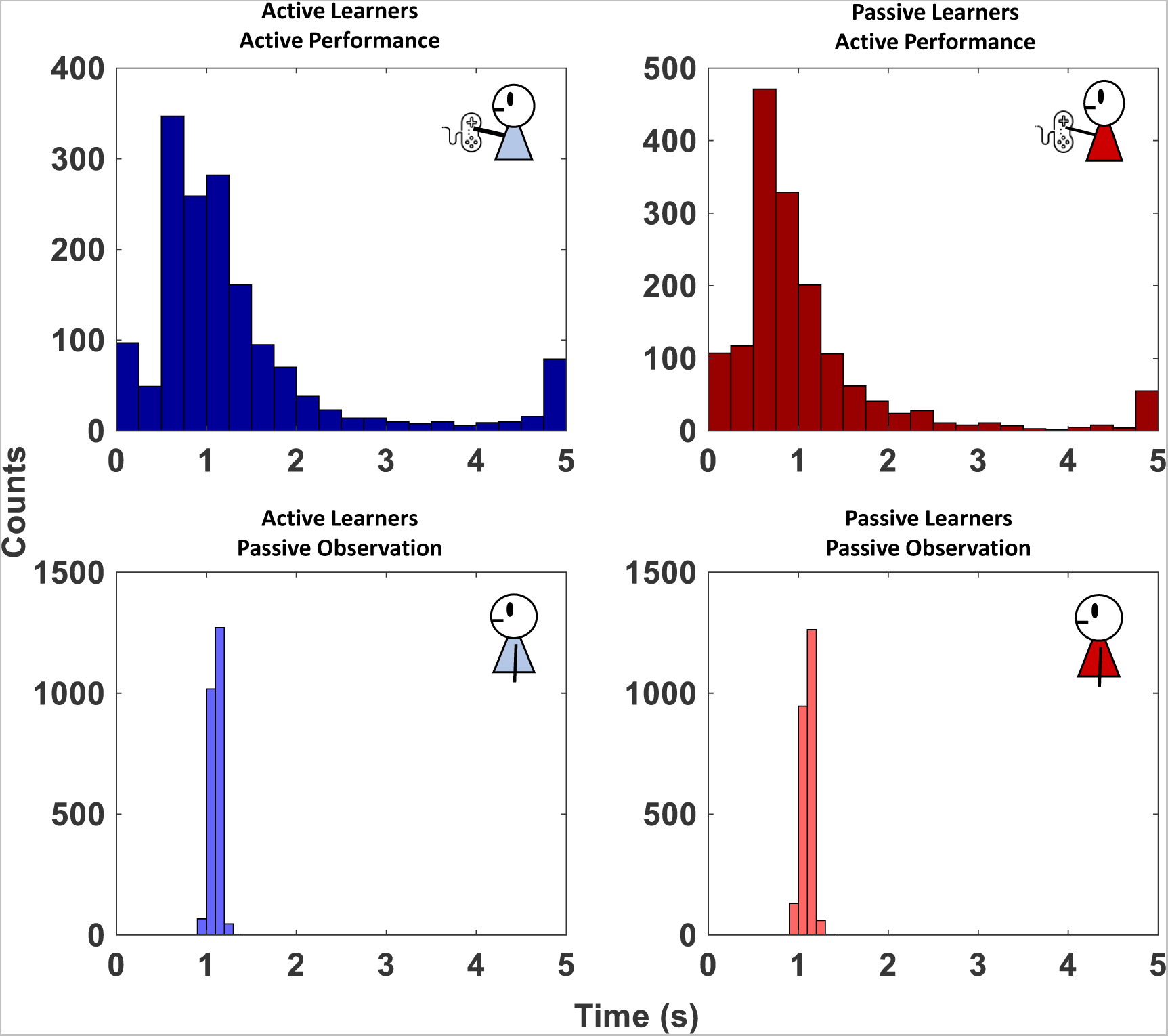
Histograms of movement durations in Active and Passive Learners in the Passive Observation and Active Performance conditions.

### Decoding of intended vs. observed movement directions

Fig. 5 shows the decoding of intended vs. observed movement directions (left vs. right) from whole-brain MEG sensor signals, in both learning groups and conditions. During Active Performance, we expected the participants to use the information from the *SC* and *CC* cues to determine the correct target location during the *CC* epoch (Fig. 5a, b, left; horizontal pale green bar), before they moved the cursor towards the target during the *MVT* epoch (Fig. 5a, b, right; horizontal grey bar). The decoders correctly classified the Left/Right hemifield of the target location decided by the Passive and Active Learners with 60-70% accuracy (*p <* 0.05; no significant differences between groups) from 170ms after the onset of the color cue (Fig. 5a, b; left, vertical grey line), and before the onset of the movement in the Active Performance condition (Fig. 5a, b; vertical grey line). Decoding accuracy increased further following movement onset, reaching about 90% over the duration of the *MVT* and *THT* epochs (Fig. 5a, b; right). This demonstrates that the MEG data are sensitive to both intended (*CC* epoch) and executed (*MVT* epoch) movement directions (left vs. right) of both the Active and Passive Learners, when they actively performed the task with overt wrist movements (Active Performance condition).

**Fig. 5.**
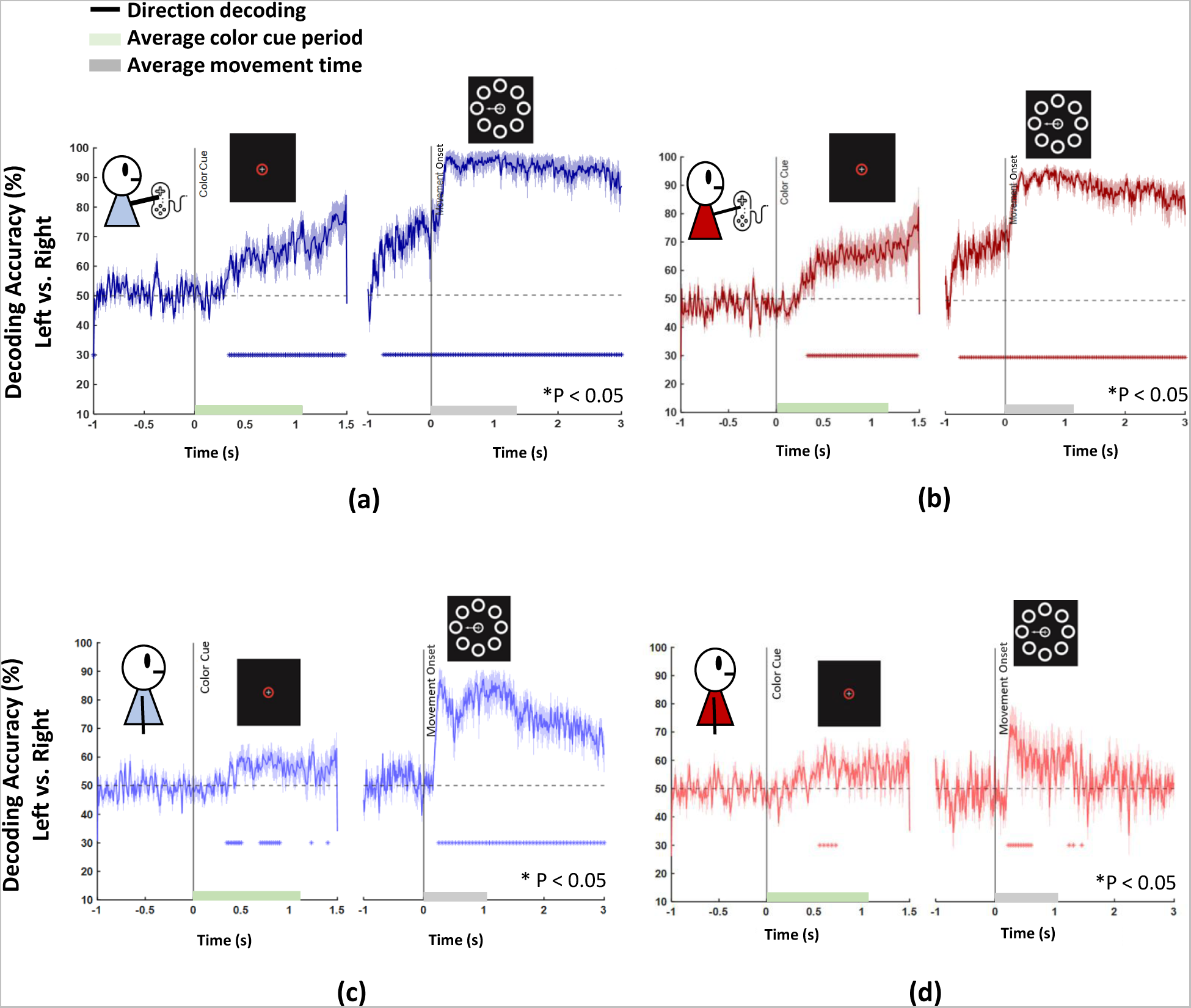
Time-resolved decoding accuracy (mean ± s.e.m.) over all MEG sensors, time-locked to color cue and movement onset, and averaged across all participants. (a) Active Learners in the Active Performance condition; (b) Passive Learners participants in the Active Performance condition; (c) Active Learners in the Passive Observation condition; (d) Passive Learners in the Passive Observation conditions. For each plot, ∗ shows times with significant decoding accuracy (signed-rank test, FDR corrected). The shaded area represents the standard error of the mean (SEM).

In the Passive Observation condition, we again expected the participants to identify the correct target after the display of the color cue (*CC* epoch), but that they would not be able to predict in which direction the computer would move the arrow (subsequent *MVT* epoch). Therefore, for the *CC* epoch of the Passive Observation condition, we sorted trials according to the correct target location based on the information conveyed by the *SC* and *CC* cues (Fig. 5c, d; left). We observed in Passive and Active Learners that the decoding accuracy reached about 60% during the *CC* epoch until the onset of computer-generated arrow movements. The decoding accuracy was not as sustained as in the Active Performance condition (Fig. 5a, b; left), but did reach above-chance levels over several time segments of the *CC* epoch (330-500 ms and 640-860ms for Active Learners, and 500-750ms for Passive Learners, *p <* 0.05). For the *MVT* epoch, we sorted the trials according to the direction of the movement of the arrow controlled by the computer, regardless of its correctness (Fig. 5c, d; right). The decoding accuracy dropped to chance level during the last 1000ms of the *CC* epoch prior to the onset of the motion of the arrow in both Passive and Active Learners. This observation is consistent with the fact that the participants were not able to predict the direction chosen by the computer, even though they presumably were aware of the correct target location, as indicated by the modest, yet significant (*p <* 0.05) decoding accuracy during the *CC* epoch in the Passive Observation condition, after sorting the trials based on the expected target location (Fig. 5c, d; left).

These decoder results confirm that the MEG data captured brain activity related to the target selected by Active and Passive Learners during the *CC* epoch in the Passive Observation condition. We were also able to decode brain activity in both Learner groups related to the target selected by the computer during the *MVT* and *THT* epochs of that condition, regardless of the participants’ prior expectations before the onset of the movement of the arrow. This latter effect was stronger and more sustained (*p <* 0.05) over time in Active (125-3000ms) than Passive (125-625ms) Learners.

We then sought to decode brain activity indicating whether the arrow moved towards the expected correct vs. unexpected incorrect direction. To that end, we sorted all Passive Observation trials into two groups for which the correct target was to the left (Fig. 6a) or the right (Fig. 6b) of the vertical midline. We sorted those two groups of “expected left” and “expected right” trials into two sub-groups, depending on whether the arrow moved toward the expected correct vs. the unexpected incorrect direction. The decoder accuracy prior to movement onset fluctuated around chance level because the participants were unable to predict whether the computer would move the arrow towards the expected correct target on the left (Fig. 6a) or the right (Fig. 6b), or towards the unexpected incorrect target location, i.e., towards the left (Fig. 6a) or towards the right (Fig. 6b). However, shortly after arrow movement onset, the decoder performance rose rapidly above chance level accuracy, discriminating between movements towards the expected correct vs. unexpected, incorrect direction. These decoding performances were statistically significant (*p <* 0.05) in Active Learners as early as 170 ms after movement onset and for the remaining duration of the *MVT* and *THT* epochs for both expected directions (Fig. 6, blue lines). The decoding accuracy was also significant albeit weaker for Passive Learners, and over a longer duration following movement onset when the expected arrow movement was towards the left direction (Fig. 6a, yellow line) than towards the right (Fig. 6b, yellow line).

**Fig. 6.**
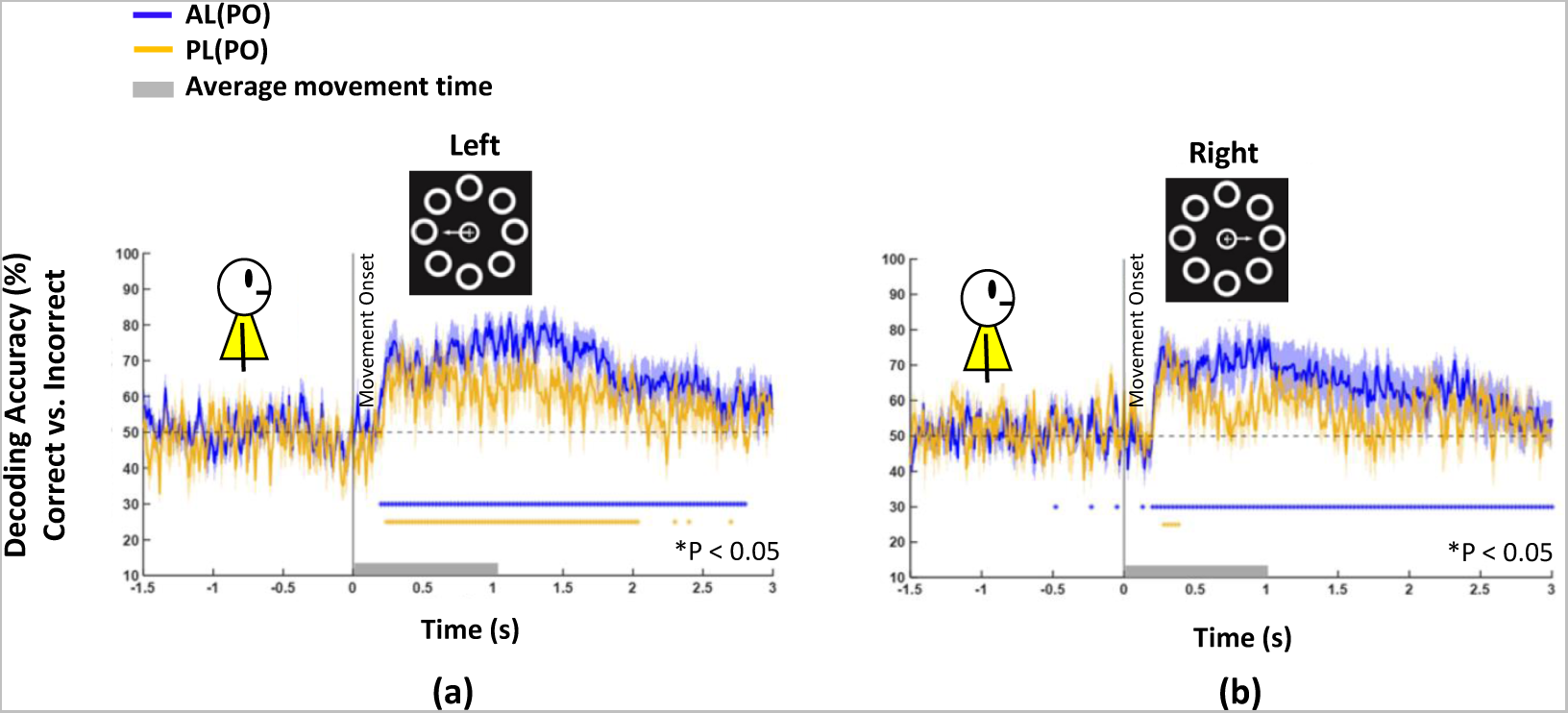
Time-resolved decoding accuracy over all MEG sensors, time-locked to movement onset for correct vs. incorrect trials (see Fig. 3) for both Active (blue) and Passive (yellow) Learners during Passive Observation. (a) Decoding accuracy of observed correct left vs. incorrect left arrow motions in trials in which participants expected the arrow to move to leftward targets. (b) Decoding accuracy of observed correct right vs. incorrect right arrow motions in trials in which participants expected the arrow to move to rightward targets. For each plot, *∗* shows times of significant decoding accuracy (signed-rank test, FDR corrected). The shaded area represents the standard error on the mean (SEM). Grey bars: duration of the arrow motion from the start position to the target (*MVT*).

### Cortical activations during Active Performance vs. Passive Observation

Fig. 7 shows the time sequence of beta-band activity in Active and Passive Learners during the *CHT, SC, CC* and *MVT* epochs of the Active Performance (Fig. 7a) and Passive Observation (Fig. 7a, b) conditions. The magnitude of beta-band activity was transformed into z-scores relative to a baseline period defined between 500 and 200ms prior to the onset of the *CHT*. The results are presented for both tasks at the same z-score scale in Fig. 7a to facilitate the relative levels of beta-band activity in the two tasks. The Passive Observation data are re-presented at a smaller z-score range in Fig. 7b to better illustrate the cortical distribution of activity changes.

**Fig. 7.**
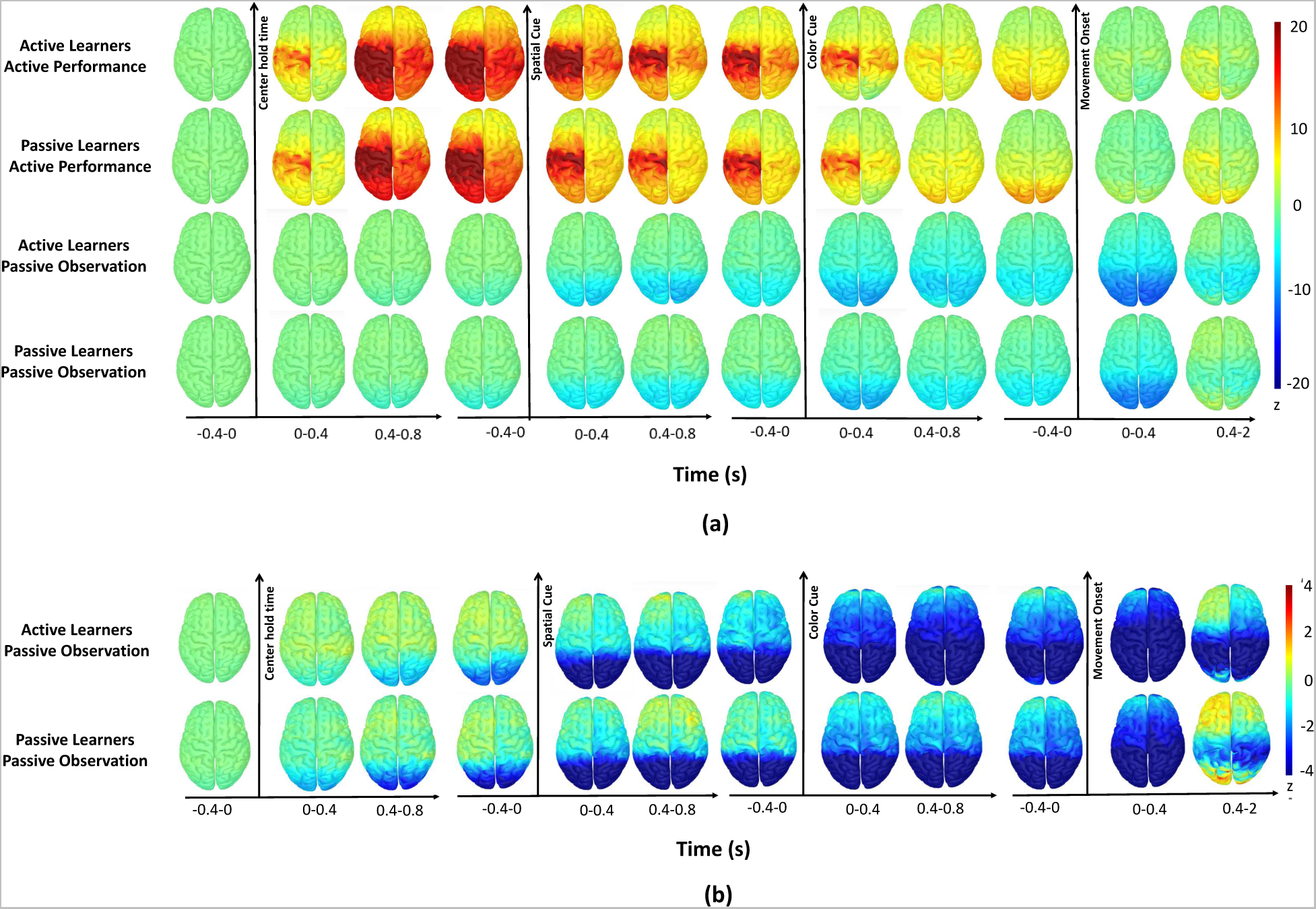
Spatiotemporal distributions of beta-band activity (15-29 Hz) (a) in the Active Performance condition for Active Learners (two top rows) and Passive Learners (two bottom rows) during *CHT, SC, CC* and *MVT* epochs. The vertical arrows indicate the respective onsets of the center-hold time, *SC*s, *CC* epochs, and movement onset, respectively. The baseline for the z-score transformation was 500 to 200 ms prior to *CHT*. (b) Same as (a) for the Passive Observation condition (note the different color scale).

Cortical beta activity was similar between Active and Passive Learners but differed depending on the Active Performance vs. Passive Observation condition (Fig. 7). Indeed, during Active Performance of both Active and Passive Learners, cortical beta activity increased from baseline during the *CHT* epoch, while the participants held the cursor in the central window, waiting for the *SC* to appear (Fig. 7a, two top rows). This increase was predominant over the motor, premotor, somatosensory, and parietal cortices of the left hemisphere, contralateral to subsequent hand movements, and extended into the occipital cortex by the end of the *CHT* epoch. We found a similar pattern in the ipsilateral hemisphere, albeit developing more slowly and of less intense amplitude. We then observed a gradual decline in beta activity in both hemispheres after the *SC* onset, with a further decrease after the presentation of the color cue *CC*, followed by increased beta activity in the occipital pole immediately prior to the expected onset of the GO cue. During the first 400ms of the *MVT* epoch, beta-band activity decreased to near-baseline levels over the bilateral motor cortex before it rebounded above baseline over the contralateral motor cortex 400-2000ms after movement onset.

In contrast, during Passive Observation, cortical beta activity did not increase during *CHT* while both the Active and Passive Learners passively observed the events driven by the computer (Fig. 7a, b). Instead, there was a progressive decrease of activity in both learner groups across subsequent epochs (Fig. 7b). We observed a suppression of activity below baseline levels first in the occipital pole, then progressing rostrally during the *SC* and *CC* epochs. This suppression was the strongest and most widespread within 400ms after the onset of the observed arrow movements. This was followed by a rebound of beta activity over the left frontal pole 400-2000ms after arrow movement onset, when the participants had to issue a decision about whether the arrow moved towards the correct target location, and count the number of correct or incorrect target choices in the current trial block. This beta rebound was also observed in the rostral sensorimotor cortex, primarily in the left hemisphere, although Active Learners did not use their right hand to manipulate the joystick during Passive Observation, and Passive Learners had not yet performed the task with the joystick.

Finally, another effect common to both Learner groups during both Active Performance and Passive Observation was the progressive suppression of beta-band activity from the onset of the *SCs* epoch until the post-movement beta rebound. This progressive suppression gradually reduced the level of beta-band activity that increased over the *CHT* during the Active Performance condition, and resulted in a decrease in beta-band activity below pre-*CHT* levels during Passive Observation.

We extracted the z-scored time course of beta-band activity in the PMd region of interest (Fig. 8c) during both Active Performance and Passive Observation in both learner groups. Our observations during Active Performance replicate those reported by (Kilavik et al., 2013) in a standard instructed-delay movement task (Fig. 8a).

**Fig. 8.**
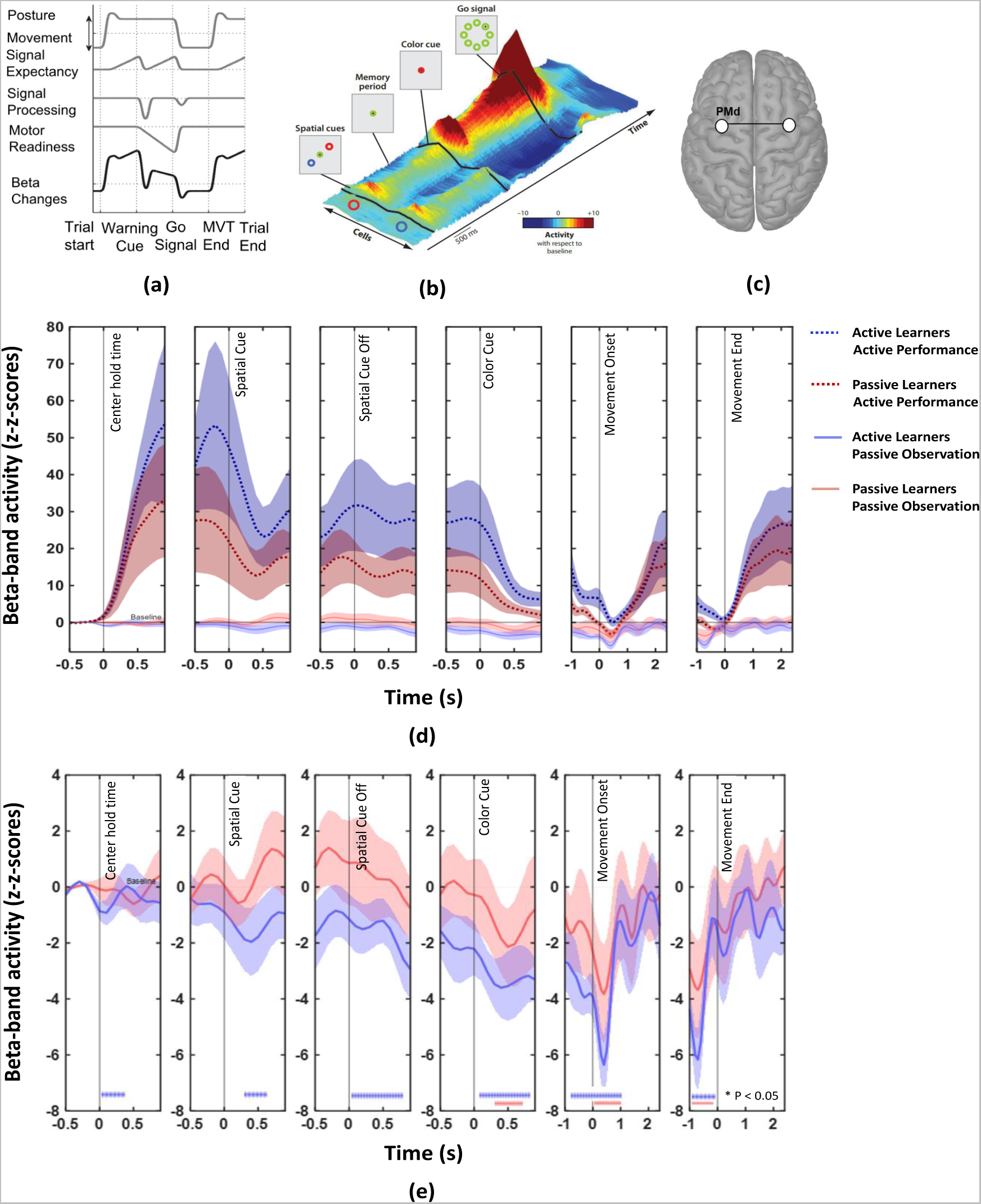
(a) Hypothesized time changes of beta activity of the premotor cortex across the task epochs of an instructed-delay motor task (modified with permission from the Experimental Neurology Kilavik et al. (2013)) (b) Dorsal premotor population activity during a 2T task, non-human primate subject (modified with permission from the Annual Review of Neuroscience Cisek and Kalaska (2005, 2010)). Two populations of PMd units that preferred two opposite potential targets were co-activated during the *SC* and *SCO* epochs. When the *CC* appeared, the activity of the units preferring the correct target increased, while the activity of the units preferring the non-selected target was suppressed. (c) Region of interest: dorsal premotor cortex (PMd). (d) Time courses of beta activity over the bilateral dorsal premotor cortex in Active and Passive Learners and in both conditions. The z-transformation was with respect to a baseline 500 to 200 ms prior to the center hold time epoch. Shaded areas show standard errors on the mean (SEM). (e) same as (d) in the Passive Observation condition with an adapted scale. *∗* indicate time segments of significant differences from baseline (*p <* 0.05; permutation tests).

We found a rapid and strong increase of premotor activity over *CHT* during Active Performance, while the participants held the cursor in the central window and awaited for the appearance of the *SC*s (Fig. 8d, dark colors). This activity then decreased substantially over the *SC* epoch as the participants were shown the locations of the two potential targets. This decrease was followed by a small rebound during the memorized *SCO* period, before decreasing again rapidly after the onset of *CC*, when the participants became able to choose the target. All these premotor effects occurred prior to movement onset, as the participants watched the visual cues and prepared to use the joystick. Premotor beta activity further decreased to near-baseline levels at the onset of joystick movements, followed by a rebound at the offset of the movement (Fig. 8d).

In contrast, during Passive Observation, we did not measure substantial changes in premotor beta-band activity during the *CHT* epoch in both Active and Passive Learners (Fig. 8d, pale colors). After the *SC* was presented, premotor activity was gradually more suppressed across the *SC, SCO* and *CC* epochs in both Learner groups. This suppression was weaker in the Passive Observation than in the Active Performance condition, even though the participants were presented the same cues and extracted the same task-relevant information from them in both conditions (Fig. 8e). Both Active and Passive Learners showed a further transient suppression of premotor activity after *SC* onset, followed by a small rebound. The onset of *CC* evoked another transient suppression of premotor beta-band activity, which decreased again sharply after the onset of arrow movements, followed by a rebound of premotor activity immediately before the arrow entered the target chosen by the computer (800ms). The main difference between the Active and Passive Learners during Passive Observation was a consistent negative offset of premotor beta-band activity initiated during the *CHT* epoch in Active Learners, which was less pronounced during the *SC* epoch in Passive Learners, followed by further progressive suppression of premotor activity for the remainder of the trial.

In summary, during Passive Observation, premotor beta activity unfolded through sequential changes that were qualitatively similar to those measured during Active Performance, albeit of lower magnitude, and without the large increase observed over the *CHT* epoch during Active Performance. Such striking similarity of premotor activity between Active Performance and Passive Observation occurred although the participants did not produce active wrist movements during Passive Observation. They simply were tasked to process the same visual cues as during Active Performance to select the correct target location and predict the appropriate direction of arrow movements. In the Passive condition, both the Active and Passive Learners issued a categorical decision about whether the arrow moved toward the correct or incorrect target.

### Differential suppression of beta activity in Active vs. Passive conditions

As noted above, one common trend we observed in both task conditions was a progressive suppression of premotor beta activity from the onset of the *SC* cue until the completion of active hand or passive arrow movements. To compare this effect directly while compensating for the large increase in beta-band activity during the *CHT* epoch of Active Performance, we defined a new pre-*SC* baseline at the end of the *CHT* epoch, 500 to 200ms prior to the onset of *SC* (Fig. 9). This baseline adjustment was meant to emphasize the change of premotor beta activity after the respective onsets of *SC* and *CC* during Active Performance in both Active and Passive Learners, and the further progressive changes in premotor activity during movement preparation between the respective onsets of *CC* and of the GO cue (Fig. 9a).

**Fig. 9.**
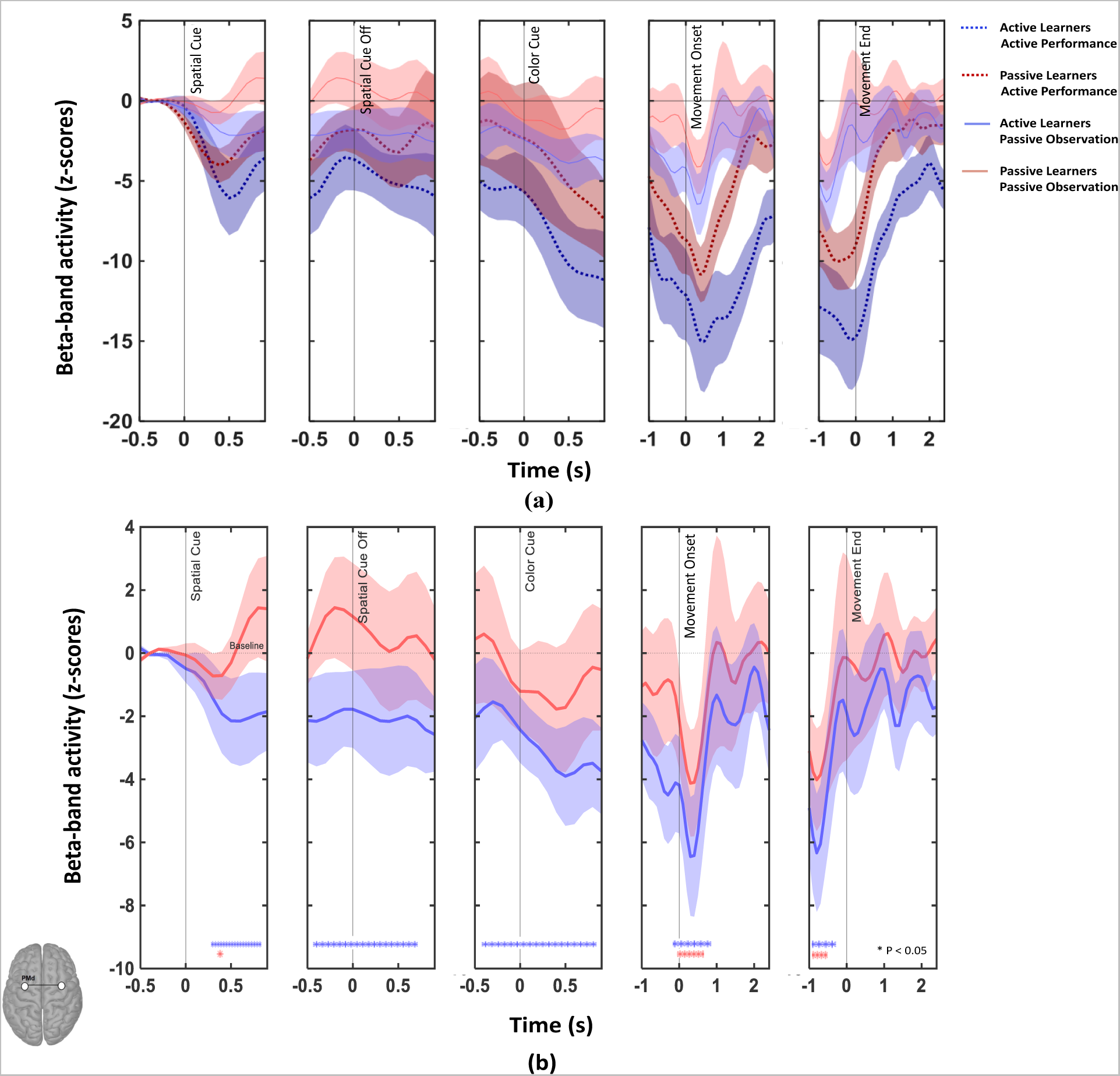
(a) Beta-band activity was z-transformed relative to a baseline defined 500 to 200 ms prior to the appearance of the spatial potential-target cue. (b) same as (a) in the Passive Observation condition (note the adapted scale to facilitate the comparison between Active and Passive Learners). The shaded area represents the standard error on the mean (SEM). *∗* along the x-axis indicate time segments of significant differences from baseline (p < 0.05; permutation tests).

This baseline adjustment emphasized the pattern of progressive decreases in premotor beta activity after the onset of *SC*s during Passive Observation in both groups of learners, albeit with reduced magnitude than during Active Performance (Fig. 9a, b). Notably, however, premotor activity was significantly (*p <* 0.05) lower than during the pre-SC baseline in Active Learners from 290ms after the onset of the SC until the rebound in the *MVT* epoch (Fig. 9b). In contrast, premotor activity did not increase significantly in Passive Learners during the SC epoch before it dropped below the pre-*SC* baseline level just before the onset of the *CC* (Fig. 9b). Premotor effects in Passive learners were significant (*p <* 0.05) at the onset of arrow motion (Fig. 9b). Furthermore, the premotor beta rebound was resolved immediately before the end of arrow movement during Passive Observation, but began at the end of the arrow/hand movement during Active Performance in both Active and Passive Learners (Fig. 9a, right). Finally, we found that with respect to movement end, the premotor rebound of activity lasted longer during Active Performance (*∼* 2000*ms* vs. *∼* 1000*ms*) (Fig. 9a, right) and was not trivially related either to greater overall beta-band activity in that condition (Fig. 8), or to the greater range of movement durations (Fig. 4).

### Differential premotor activity during passive observation of correct versus incorrect task performance

Fig. 10 illustrates the time course of premotor beta activity during Passive Observation in Passive (Fig. 10a) and Active Learners (Fig. 10b), with respect to a baseline reference defined from 1000 to 200ms prior to the GO cue and arrow movement onset. Premotor activity was transiently suppressed at the onset of arrow movements, prior to a marked rebound (500ms after movement onset), in both learner groups. In Passive Learners during Passive Observation, we found stronger rebounds of premotor activity in incorrect than in correct trials during arrow movements (Fig. 10a). This difference was significant 1100-1300 ms after movement onset, and 1400-1600ms after movement end (*p <* 0.05). In contrast, also during Passive Observation, we noted stronger rebounds of premotor activity in correct than incorrect trials in Active Learners (Fig. 10b). This difference was significant 600-700ms after movement onset, 800-400 ms before movement ended, and 1700-1800ms after movement ended (*p <* 0.05). In both cases, the effect was most pronounced in the higher range of the beta band (22-29Hz). In summary, we found evidence of differential expressions of premotor activity during Passive Observation depending on correct vs. incorrect arrow movements.

**Fig. 10.**
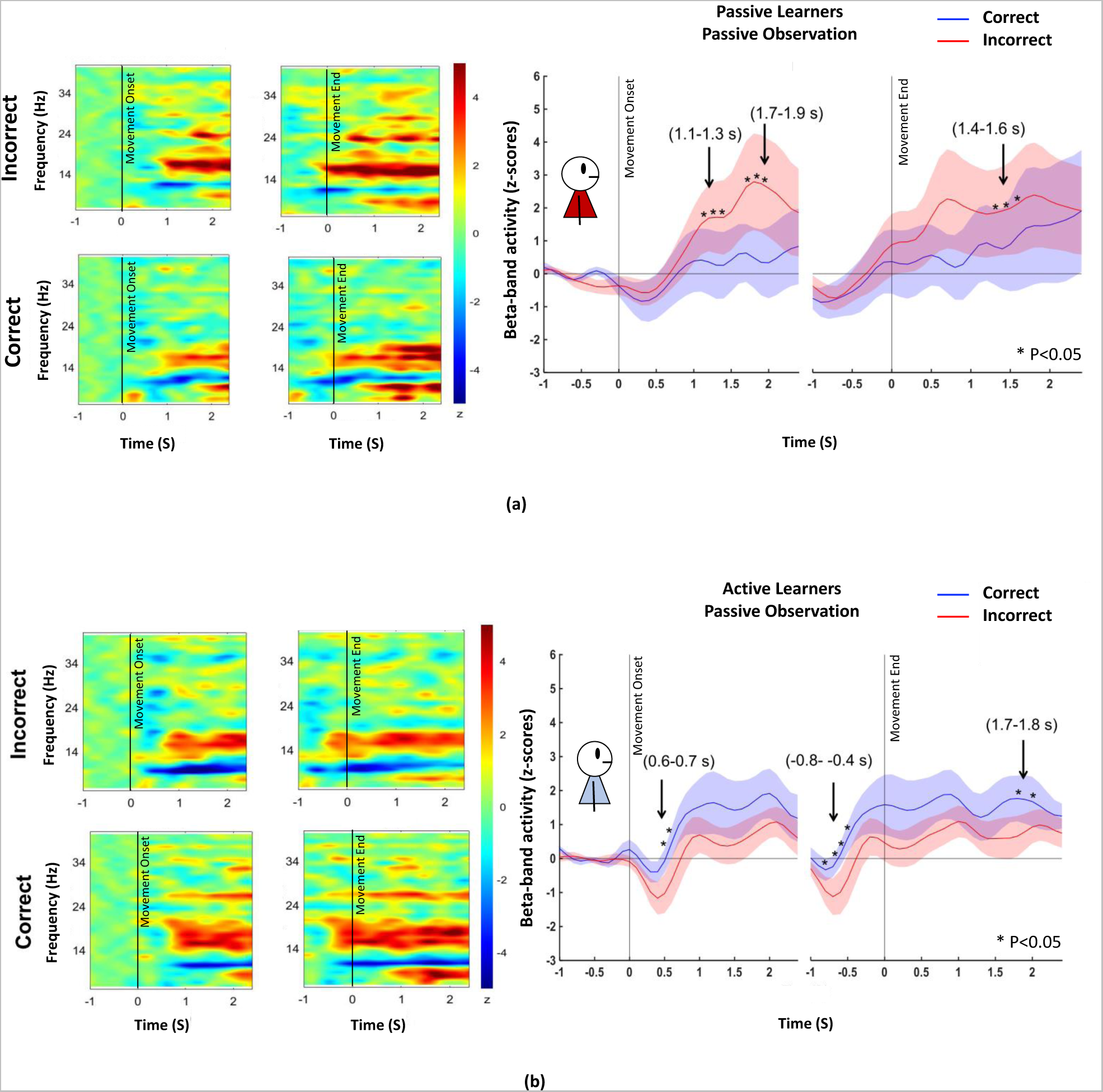
Grand average spectrogram for correct and incorrect trials in the Passive Observation condition, for both groups of learners. The baseline of the z-transformation was taken from 1000 to 200 ms prior to movement onset. (a) Passive Learners showed a stronger beta rebound in incorrect trials of the Passive Observation condition, with an opposite pattern in Active Learners (b). The shaded area represents the standard error of the mean (SEM). *∗* shows time segments of significant differences between observed correct and incorrect responses (pair-wise t-test, *p <* 0.05).

## Discussion

We adapted for human participants and non-invasive brain imaging a “Two-Target” (2T) task originally developed for non-human primates (Cisek and Kalaska, 2004, 2005). We measured brain activations related to 2T stimulus interpretation, outcome prediction and error processing during the observation of a sequence of non-biological stimuli that follow an arbitrary but simple stimulus-response association rule. The aim was to determine whether motor training with overt hand movements is a prerequisite for the involvement of human premotor cortex during the covert cognitive appraisal of the arbitrary 2T visual stimulus-response events.

We measured changes in beta-band activity with respect to reference baselines using time-resolved MEG source mapping across the whole cortex and with specific investigations in the dorsal premotor cortex. We compared effects when the participants performed the 2T task actively using wrist movements to report their target choice (Active Performance), or passively while they applied the same color-location conjunction rule to make a categorical decision about whether a computer performed the task correctly (Passive Observation). We identified differential cortical activity in participants who either learned the task via trial-and-error during active task performance (Active Learners), or by passive observation of the trial outcomes when a computer performed trials correctly or incorrectly (Passive Learners). Finally, we investigated whether premotor activity during Passive Observation distinguished between the correct and incorrect performance of the task by a computer.

Although beta-band effects have been extensively documented in a variety of motor tasks, their functional relevance so far has been uncertain (Sanes and Donoghue, 1993; Donoghue et al., 1998; Pfurtscheller, 1992; Pfurtscheller et al., 1996; Pfurtscheller and Da Silva, 1999; Doyle et al., 2005; Alegre et al., 2006; Jurkiewicz et al., 2006; Pesaran et al., 2008; Van Wijk et al., 2009; Donner et al., 2009; Engel and Fries, 2010; Saleh et al., 2010; Haegens et al., 2011; Kilavik et al., 2012, 2013; Cheyne, 2013; Khanna and Carmena, 2017; Rule et al., 2017, 2018; Little et al., 2019; Barone and Rossiter, 2021; van Helvert et al., 2021). Beta-band activity during active motor tasks has been attributed such diverse roles as an “idling rhythm” related to movement inhibition or maintenance of the current functional state of motor circuits, attentional processes, expectation of imminent task-salient sensory events and movement onsets, response selection, and other sensorimotor decision-making mechanisms underlying voluntary motor control. However, beta suppressions and rebounds also arise, albeit typically with smaller amplitudes, in task conditions that do not involve overt motor performance, such as during covert motor imagery (Schnitzler et al., 1997; Pfurtscheller and Neuper, 1997; Pfurtscheller and Da Silva, 1999; McFarland et al., 2000; Pfurtscheller et al., 2005; De Lange et al., 2008; Pfurtscheller and Solis-Escalante, 2009; Solis-Escalante et al., 2012; Nakagawa et al., 2011; Brinkman et al., 2014) as well as during passive observation of actions performed by a third person (Babiloni et al., 2002, 2016; Järveläinen et al., 2004; Muthukumaraswamy and Johnson, 2004; Muthukumaraswamy et al., 2006; Koelewijn et al., 2008; Orgs et al., 2008; Kilner et al., 2009b; Press et al., 2011; Cannon et al., 2014; Meyer et al., 2016). The generation and modulation of beta-band activity in those non-motor tasks presents further challenges for understanding its causal origin and functional roles.

### Premotor selection and planning of voluntary movements

One advantage of this MEG study is that the behavioral psychophysics and single-neuron responses in PMd and M1 are well documented in NHP studies of the 2T task (Cisek and Kalaska, 2004, 2005; Coallier and Kalaska, 2014; Coallier et al., 2015; Wang et al., 2019). Our present data show that in both Active and Passive Learners, cortical activity during Active Performance is similar between the two groups of learners (Fig. 7a, top two rows). We found a rapid increase of beta activity during the *CHT* epoch relative to a pre-*CHT* baseline, initiated in sensorimotor cortices, and spreading rapidly to other cortical regions. Beta activity then decreased progressively in a stepwise fashion during the *SC, SCO,* and *CC* epochs, returning to pre-*CHT* baseline levels at the onset of wrist movements, followed by a rapid rebound beginning at the end of active wrist movements. These effects were bilateral but stronger in the hemisphere contralateral to the hand used to perform the task. Moreover, we showed there was enough differential information in the patterns of cortical MEG signals to decode the direction towards which both learner groups intended to make wrist movements during Active Performance as soon as the *CC* was displayed (Fig. 5).

During Active Performance, we found temporal patterns of premotor activity (Fig. 8d) similar to those observed in studies of reaction-time and instructed-delay motor tasks (Sanes and Donoghue, 1993; Donoghue et al., 1998; Doyle et al., 2005; Alegre et al., 2006; Zhang et al., 2008; Van Wijk et al., 2009; Saleh et al., 2010; Park et al., 2013; Torrecillos et al., 2015; Tzagarakis et al., 2010, 2015; Kilavik et al., 2012, 2013; Brinkman et al., 2014; Rule et al., 2017, 2018; van Helvert et al., 2021). Of particular interest, (Saleh et al., 2010) also used a color-location conjunction rule in an 8-target variant of the 2T task (“color-association” task). They recorded local field potentials with a 96-microelectrode array chronically implanted in the M1 hand region of a human tetraplegic patient. In each trial, the patient observed 8 color-coded targets and 5 different sequential central color cues that each corresponded to the color of one of the 8 potential targets. However, only the second or fourth cues in the sequence defined the correct target in different trials. The participant made a chin movement to move a cursor to the target at the onset of a GO cue after the fifth color cue. Aligned with our present findings, their data showed a large increase of beta-band activity during the initial central starting-hold period before the onset of the first color cue, followed by a transient increase before and a transient suppression after the appearance of the first to fourth color cues: a progressive step-wise decrease in beta-band activity with each successive cue, and a strong decrease after the onset of the second or fourth cues that defined the correct target in each trial. This beta suppression was sustained until the GO cue (no effect after the GO cue was reported in their study). This temporal profile strikingly parallels our present results during Active Performance.

Another relevant finding of (Saleh et al., 2010) was that the time course of beta-band activity was altered when the five color cues of the color-association task were replaced with five sequential monochromatic circles that appeared at five different sequential potential target locations (“spatial-location task”; Saleh et al. (2010)). In contrast to the color-association task, they found that the initial increase in beta activity did not occur until after the first target cue appeared in the spatial-location task. Furthermore, the transient decrease in beta activity after the appearance of each color cue in the color-association task did not occur after the appearance of each of the target cues in the spatial-location task. We interpret each transient decrease in beta-band activity after the appearance of the color cues in their color-association task as marking when the participants applied the color-location conjunction rule to locate the target that corresponded to the current color cue. That information-processing step was not required in the spatial-location task because each monochromatic cue signalled explicitly the location of the corresponding target. Therefore, the differences in the beta-band time profile reflected an alteration of the information-processing mechanisms used in each respective task (Saleh et al., 2010).

The prominent initial increase of beta activity we report during the initial hold period of the *CHT* epoch during Active Performance replicates several prior studies (Sanes and Donoghue, 1993; Donoghue et al., 1998; Van Wijk et al., 2009; Rubino et al., 2006; Saleh et al., 2010; Takahashi et al., 2011; Kilavik et al., 2012, 2013; Rule et al., 2017). In each of these studies, participants performed a distinct motor act (other than gaze fixation) to initiate their active engagement in each trial and intended to perform a further motor act in response to the instructional cues that appeared during the trial. It is possible, therefore, that an increase in beta activity at the start of a trial may be modest or absent in tasks where the participants do not perform a motor action to initiate a trial before the appearance of the first instructional cue, such as during the present Passive Observation condition. Furthermore, most of the studies that did not report a beta-band increase at the start of trials, the baseline of reference for beta-band activity changes was defined immediately before the onset of the first instructional cue. As a result, any increase in beta activity that might have occurred prior to that time window in each trial was not quantified and reported. We demonstrate this effect using a baseline time window immediately prior to *CHT* (Fig. 8), as opposed to using a baseline near the end of *CHT* before the onset of the *SC*s (Fig. 9).

Increased beta activity has often been associated with an “idling” state during, e.g., the maintenance of a posture or inhibition of movement (Pfurtscheller, 1992; Pfurtscheller and Neuper, 1994; Pfurtscheller et al., 1996; Pfurtscheller and Da Silva, 1999; Alegre et al., 2006; Gilbertson et al., 2005; Solis-Escalante et al., 2012; Kilavik et al., 2013). In the 2T task, however, the premotor circuits are not idle during the instructed-delay periods, but are rather expected to participate in the processing of the information conveyed be each cue (Cisek and Kalaska, 2004, 2005; Coallier and Kalaska, 2014; Coallier et al., 2015; Wang et al., 2019). We also measured large increases in beta activity during CHT in both Active and Passive Learners, followed by progressive decreases at the onset of each new instructed-delay epoch while the participants maintained a constant wrist posture until the onset of the GO cue. Moreover, the need to withhold movement was greater during the *CC* epoch when the participants had already decided which wrist movement they were about to produce, but were instructed to wait for the GO cue; yet beta-band activity returned back to near pre-CHT baseline levels during that epoch. Finally, post-movement beta rebounds have been associated with the notion of active inhibition of neural networks after movement execution (Solis-Escalante et al., 2012) or a resetting of cortical circuits to their pre-movement state following a motor performance (Zhang et al., 2008). However, we found similar levels of beta activity during the post-movement rebound while the participants held the cursor at the chosen target location during Active Performance as during the *SCO* epoch while they were holding the cursor at the central start position and awaiting the appearance of the *CC* to choose the correct target (Fig. 8d). Importantly, NHP data show that the levels and patterns of premotor neural activity are distinct between these two epochs, although beta-band activity remains relatively stable(Cisek and Kalaska, 2004, 2005; Coallier et al., 2015; Wang et al., 2019). During the *SC* and *SCO* epochs, two sub-populations of PMd neurons increase their activity simultaneously to represent the two potential movements to the two targets (Cisek and Kalaska, 2004, 2005; Coallier et al., 2015; Wang et al., 2019). In contrast, PMd single-neuron activity during beta activity rebounds over the target-hold period at the end of movements is a complex mixture of single-unit discharge patterns related to the maintenance of the current posture of the arm and to the anticipation of the return movement to restore the cursor at the central position to initiate the next trial (Crammond and Kalaska, 1996, 2000; Cisek and Kalaska, 2005; Coallier et al., 2015; Wang et al., 2019). In sum, these complex single-neuron activity patterns do not seem to reflect either an active inhibition of neural circuits or a generic resetting of cortical circuits during the post-movement beta rebound period.

In contrast to idling, postural maintenance or circuit resetting, the changes in beta activity at each transition between trial epochs during Active Performance occurred whenever there was a distinct change in the information processed during each epoch. The large increase of beta activity during the initial *CHT* epoch occurred after the participants positioned the cursor at the central start position and actively maintained the cursor at the central target location while waiting for the onset of the first instructional cue (*SC*). We then observed an abrupt decrease of beta activity at the onset of *SC*, which informed the participants of the color-location conjunction of the two targets for the current trial. This increase was followed by a rebound of beta activity immediately before the offset of *SC*. Beta activity then remained stable over the duration of the *SCO* epoch, during which time the participants retained in memory the specific color-location conjunction corresponding to the two potential targets. At the onset of the *CC*, the participants chose the correct target by recalling the memorized target color-location conjunction for the current trial and matching it to the color of the *CC*, before they prepared for the associated wrist movement. This was accompanied by a second abrupt reduction of beta-band activity that was sustained for the duration of the *CC* epoch, during which the participants withheld their chosen movement. The sharp decrease in beta activity after the onset of the *CC* also marked a transition from a period of uncertainty about the action to perform during the *SC/SCO* epochs to one of certainty about the chosen action during the *CC* epoch, consistent with previous findings of the effect of response certainty on beta-band activity in the sensorimotor cortex (Doyle et al., 2005; Donner et al., 2009; Tzagarakis et al., 2010, 2015; Grent-’t Jong et al., 2014; van Helvert et al., 2021). Finally, the onset of the GO cue resulted in a further transient suppression of beta activity during the initiation and execution of the chosen wrist movement, followed by an abrupt rebound of beta activity after the arrow entered the target (Fig. 8d). These observations mirror the changes in premotor neural activity patterns during each trial epoch while NHPs performed reaching movements in the same 2T task (Cisek and Kalaska, 2004, 2005; Coallier et al., 2015). The first large increase in beta activity during the *CHT* epoch occurred when the subjects became actively engaged in the trial by positioning the cursor in the central start window. Interestingly, this trial segment was not accompanied by a significant change in PMd tonic discharge rates (Crammond and Kalaska, 1996, 2000; Cisek and Kalaska, 2005; Coallier et al., 2015). The first decrease in beta activity during the *SC* and *SCO* epochs occurs when two sub-populations of PMd neurons are coactivated to represent the two potential reaching movements in the current trial. The second beta decrease occurs after the onset of *CC*, when the subjects need to choose their final target and withhold the corresponding motor response until the onset of the GO cue. This resulted in a large increase in the activity of the neural sub-population that preferred the chosen reach movement and a strong suppression of the PMd neurons that preferred the other, rejected, target. Finally, the suppression of beta activity during movements corresponds to when the PMd population generates a strong movement-related burst of activity. The post-movement beta rebound occurs when mean PMd neural activity decreases substantially after movement ends but different single neurons often show correlations with either the maintenance of posture over the chosen target or the anticipated return movement back to the central start position to initiate the next trial (Crammond and Kalaska, 1996, 2000; Cisek and Kalaska, 2005; Coallier et al., 2015).

Finally, recent computational models propose new functional interpretations of beta-band activity in motor tasks. According to these models, primary motor and premotor neural circuits are conceived as passing through a succession of activation states in the course of instructed-delay tasks (Churchland et al., 2010, 2012; Shenoy et al., 2013; Kaufman et al., 2014; Cunningham and Yu, 2014; Elsayed et al., 2016; Gallego et al., 2017, 2018; Kalaska, 2019; Vyas et al., 2020; Duncker and Sahani, 2021; Thura et al., 2022; Meirhaeghe et al., 2023). These states include “output-null” states during which neural circuits select and prepare a movement without generating muscle activity during instructed-delay periods. This period then transitions to an “output-potent” state during which the neural circuits can generate muscle activity to produce overt movements after the onset of the GO cue. Synaptic interactions and discharge correlations within the network may enable such transitions and are marked by oscillatory signal components measurable at the mesoscopic scale, such as with MEG source imaging as in the present study (Sanes and Donoghue, 1993; Murthy and Fetz, 1996a,b; Donoghue et al., 1998; Pfurtscheller and Da Silva, 1999; Pesaran et al., 2008; Denker et al., 2007, 2011).

In sum, we propose that during Active Performance, beta band activity in the premotor cortex reflects a sequence of information-processing events for selecting and executing a motor output that matches visual instructional cues (Doyle et al., 2005; Alegre et al., 2006; Zhang et al., 2008; Saleh et al., 2010; Kilavik et al., 2013; Park et al., 2013; Brinkman et al., 2014; Little et al., 2019), and is associated with dynamic patterns of single-unit discharges (Crammond and Kalaska, 1996, 2000; Cisek and Kalaska, 2005; Coallier et al., 2015; Wang et al., 2019). These discharges result from the computational state of the underlying neural circuits (Churchland et al., 2010, 2012; Shenoy et al., 2013; Kaufman et al., 2014; Cunningham and Yu, 2014; Elsayed et al., 2016; Gallego et al., 2017, 2018; Kalaska, 2019; Vyas et al., 2020; Duncker and Sahani, 2021; Thura et al., 2022; Meirhaeghe et al., 2023), which are reflected, in turn, in the global oscillatory state of the network, including in the beta frequency range (Sanes and Donoghue, 1993; Murthy and Fetz, 1996a; Donoghue et al., 1998; Pfurtscheller and Da Silva, 1999; Pesaran et al., 2008; Denker et al., 2007, 2011; Rule et al., 2017, 2018).

### Premotor activity during the passive observation of non-biological visual events

The present study also assessed the involvement of the human premotor cortex in categorical decisions based on arbitrary non-biological sensory events, and that do not lead to overt motor responses. We designed the Passive Observation condition of the 2T task during which the participants were presented the same visual events as in the Active Performance condition, and where they were tasked to decide whether a computer moved the arrow towards the correct target location, and keep the count of the number of correct or incorrect observed outcomes.

We found using decoding approaches in the Passive Observation conditions in both Learner groups that whole-brain MEG signals reflect both the expected direction of the arrow motion after presentation of the color cue, and the actual direction of arrow motion. We note that the decoders performed with greater accuracy with Active than with Passive Learners (Figs. 5, 6).

The first novel finding is that the strong increase in sensorimotor beta activity above pre-*CHT* baseline during the instructed-delay periods that occurred in the Active Performance condition did not occur in the Passive Observation task. The absence of the equivalent response in the Passive Observation condition cannot be attributed to a lack of attention (Saleh et al., 2010), because the participants were tasked in that condition to register the visual cues to identify the correct target before making a covert categorical decision about whether the computer’s target choice respected or violated the color-location conjunction rule. They were also required to count the number of correct or incorrect trials performed by the computer. This latter covert cognitive act of counting alone requires substantial attentional resources (Wilder et al., 2009). Alternatively, the surge of beta activity in the Active Performance condition is likely to reflect the active engagement of the premotor cortex and other sensorimotor cortical regions in a series of information-processing events required to select and execute overt report movements of the hand to the correctly colored target in response to the sensory information from the environment. During Passive Observation, the participants did not hold nor use a joystick and did not use the visual cues to select a hand movement to report their target choice. We suggest that there was no increase in premotor beta activity during the initial part of each trial in the Passive Observation condition because no overt motor actions were required in that condition to start each trial and to report their categorical decision.

The second novel finding consists of the temporally structured fluctuations in premotor beta activity during Passive Observation, expressed as a series of progressive decreases in activity relative to baseline during the instructed-delay periods, and synchronized with the succession of trial events. The dynamical profile of these fluctuations from the start of the SC epoch to the end of the trial is qualitatively similar to those of the Active Performance condition but of lesser magnitude (Figs. 7, 8, 9). We noted the decrease of beta activity was initiated in occipital regions during the *CHT* epoch (Pfurtscheller, 1992; Little et al., 2019), before spreading over rostral cortical regions during the *SC, SCO* and *CC* epochs (Fig. 7). We found that the decrease of beta activity was maximal when the computer initiated the movement of the arrow toward one of the two possible targets. Once the movement was completed and as the participants assessed the correctness of the computer performance, we noted a rebound of beta activity over contralateral frontal and dorsal premotor regions (Fig. 7, Figs. 8, 9). Overall, these data show that the human premotor cortex is activated during the observation of arbitrary non-biological visual events that do not display familiar goal-directed actions of actors, avatars, robotic devices, or tools but only require a categorical decision based on the observed events (Schubotz and von Cramon, 2001, 2002a,b; Wolfensteller et al., 2007; Engel et al., 2008b,a; Wang et al., 2015).

The suppression of beta activity during Passive Observation of instructed-delay periods followed by a rebound are consistent with previous studies of biological action observation (Babiloni et al., 2002, 2016; Järveläinen et al., 2004; Muthukumaraswamy and Johnson, 2004; Muthukumaraswamy et al., 2006; Orgs et al., 2008; Koelewijn et al., 2008; Kilner et al., 2009b; Press et al., 2011; Cannon et al., 2014; Meyer et al., 2016) or covert mental simulation of movements (Schnitzler et al., 1997; Pfurtscheller and Neuper, 1997; McFarland et al., 2000; Pfurtscheller et al., 2005, 2006; De Lange et al., 2008; Pfurtscheller and Solis-Escalante, 2009; Solis-Escalante et al., 2012; Nakagawa et al., 2011; Brinkman et al., 2014).

We also aimed to determine whether the human premotor cortex is involved in the interpretation of arbitrary sensory events only after learning the association rule between those events and specific motor actions, or could be activated without any prior motor association. This question aimed to clarify the implication of the human premotor in embodied decision-making.

We report activity in the premotor cortex of Active Learners during Passive Observation after they had learned the 2T task using hand movements to report their target choice decision in each trial. This is reminiscent of (Cisek and Kalaska, 2004), who found that PMd neurons expressed in non-human primates similar responses during their active performance of the 2T task and while they passively watched stimulus events unfold on a computer monitor while an experimenter performed the 2T task out of sight. Consistent with Cisek and Kalaska (2004), we may therefore argue that during Passive Observation, Active Learners simulate mentally the learned motor responses associated with visual cues to predict the correct target location and where the computer should move the arrow (Schnitzler et al., 1997; Muthukumaraswamy et al., 2006; Caetano et al., 2007; Pfurtscheller and Da Silva, 1999; McFarland et al., 2000; De Lange et al., 2008; Pfurtscheller and Solis-Escalante, 2009; Nakagawa et al., 2011; Solis-Escalante et al., 2012; Brinkman et al., 2014). Such embodied internal representations may therefore enable Active Learners to make categorical decisions about the correctness of task execution by the computer.

The third novel finding of the present study is that the premotor cortex of Passive learners also expresses during Passive Observation a progressive suppression of beta activity during Passive Observation, after they have learned the color-location conjunction rule by passive observation of the stimulus events and knowledge-of-results feedback on the correctness of outcomes. Critically, and unlike Active Learners, Passive Learners did not learn to associate the stimulus events with specific hand movements. The premotor cortex has long been attributed a role in the processing of visual instructional cues that guide movement selection according to specific stimulus-response association rules (Cisek et al., 2003; Wang et al., 2019; Coallier and Kalaska, 2014; Cisek and Kalaska, 2004; Crammond and Kalaska, 1994; Halsband and Passingham, 1985; Mitz et al., 1991; Muhammad et al., 2006; Rossi-Pool et al., 2016, 2017; Wallis and Miller, 2003; Wise et al., 1996, 1997; Hoshi and Tanji, 2002, 2006; Nakayama et al., 2008; Yamagata et al., 2009). Our findings suggest that the premotor circuits of human Passive Learners contribute to the interpretation and assessment of whether non-biological visual events respect a stimulus-response association rule, even though these participants are not required to issue overt motor responses. During Passive Observation, the modulation of post-*CHT* beta activity was weaker in Passive than in Active Learners. We suggest this may be due to distinct inter-individual variability of beta-band modulations between the two groups. Indeed, beta-band modulations were also weaker in Passive Learners in the Active Performance condition (Fig. 8d). Alternatively, this difference might have resulted from the order in which the participants learned the tasks. Active Learners experienced the Passive Observation condition during their second MEG recording session after having actively performed the same task during their first visit. In contrast, Passive Learners experienced the Passive Observation condition during their first visit, immediately after learning the stimulus-response rule that determined correct trial outcomes. The stronger beta band modulations observed in Active Learners might therefore reflect a consolidation or practice effect because they performed the Passive Observation task in their second recording session.

Alternatively, the enhanced effects of beta-band modulations and greater decoding accuracy from Active Learners in the Passive Observation condition could be a manifestion of neurophysiological mechanisms of embodied decision-making. Because of their prior Active Performance motor experience with the task, Active Learners in the Passive Observation condition were able to interpret the visual stimuli in a “motor space” of potential hand movements in part by covert recall and simulation of the associated wrist movements they had learned previously (Cisek, 2006; Cisek and Kalaska, 2004, 2010; Gold and Shadlen, 2007). In contrast, Passive Learners experienced the Passive Observation condition without the prior opportunity to relate the visual stimuli to specific hand movements through active practice. We acknowledge that this possible explanation remains speculative at this point. Furthermore, while Passive Learners did not use the joystick before experiencing the Passive Observation condition, all participants had years of experience using a computer mouse, trackpad or joystick to interact with computers. For this reason, we intentionally used a growing arrow as a unfamiliar user-computer interface report mechanism. Nevertheless, we cannot reject the possibility that Passive Learners may have engaged in some form of covert motor imagery related to the motion of the arrow while in the Passive Observation condition.

Regardless of the possible involvement of mental motor simulation, the present data demonstrate that during Passive Observation, the premotor cortex of human Passive Learners is engaged while extracting information from visual stimuli and making a categorical decision about whether the observed events respect or violate a color-location conjunction rule previously learned by passively observing a computer’s performance of the task.

To address the possible confounding factor of the order of learning and task performance, both Active and Passive Learners would ideally need to be re-tested in the Passive Observation condition over a third visit, after both groups performed/observed the 2T task in the Active and Passive conditions. We would then expect the group differences in premotor beta activity to be reduced, if a form of embodied decision-making involving simulations of the specific learned wrist movements had contributed to the greater beta-band modulations observed in Active Learners during Passive Observation.

Finally, the fourth novel finding of this study is the differences in the amplitude of beta rebound activity in response to correct and incorrect target choices made by a computer in the Passive Observation task, in both Active and Passive Learners. Previous fMRI and EEG beta-band effects have been reported when participants observe human actors perform familiar and ecologically natural arm and hand movements correctly or incorrectly (Manthey et al., 2003; Meyer et al., 2016). Consistent with the present findings, (Koelewijn et al., 2008) also found differences in MEG beta-band rebound amplitude when participants observed a human actor chose a correct or incorrect hand movement or target button in response to a complex visual instructional cue and an arbitrary stimulus-response association rule.

In contrast, (Wang et al., 2015) found no difference in EEG alpha-band (8-12 Hz) power during the observation of correct versus incorrect action choices made by a computer in a symbolic Flanker-like task that used non-biological visual stimuli and no observable actors. Finally, a series of studies reported changes in fMRI BOLD signal levels in several cortical regions when participants were tasked to detect violations of the sequential structure of a series of arbitrary non-biological stimuli (Schubotz and von Cramon, 2001, 2002a,b; Schubotz and Von Cramon, 2004; Wolfensteller et al., 2007). However, these authors did not report differential brain responses to stimuli that respected or violated the sequential structure, nor effects related with the participants’ categorical decisions.

From this literature review, we believe the present study is first to report differential responses of the human premotor cortex during the observation of arbitrary events in the context of a stimulus-response association rule, without the involvement of biological motion nor actors in plain sight. We report effects of the Passive Observation of the correctness of cursor arrow motions on the amplitude of premotor beta rebounds, consistent with previous observations of error-dependent MEG beta signalling from (Koelewijn et al., 2008). We found a marked difference between the effects detected in Active and Passive Learners. Whether this effect reflects uneven inter-individual variability, is caused by different task orders or practice between the two groups, or by the actual engagement of embodiment mechanisms in Active Learners but not in Passive Learners remains to be answered. Furthermore, the outcome-related differences were only statistically significant over brief time segments of the beta rebound in both Learner groups. This suggests an outcome-related effect on human premotor beta-band activity during passive observation of a computer task performance, thereby providing initial evidence for the involvement of the human motor system in the understanding and interpretation of non-biological visual events.

Another current limiting factor may be that in the participants were tasked to count the number of correct or incorrect trial outcomes in different trial blocks (van Schie et al., 2004; Koelewijn et al., 2008). By doing so, we aimed to avoid conflating brain activity related to categorical decision making with brain activity associated with the selection of a subsequent overt motor action. However, this design strategy imposed that we could not index the participants’ categorical decisions on a trial-by-trial basis. Instead, we were only able to sort trials based on the computer’s correct versus incorrect target choices. We argue that it is reasonable to assume that the participants’ correct decisions in the Passive Observation condition were at ceiling, because their performances was 100% correct in the Active Performance condition. Nevertheless, we acknowledge that some uncertainty remains about the possible confound from sorting trials according to the computer’s target choices rather than the participants’ actual report of their trial-to-trial categorical decisions.

Finally, analyses were largely based on multi-trial averaged beta-band signals. However, single-trial analyses have shown that beta-band activity usually occurs in transient bursts at highly variable times within trial epochs and across trials (Murthy and Fetz, 1996a; Denker et al., 2007; Rule et al., 2017, 2018; Little et al., 2019; Zich et al., 2023). Nevertheless, the frequency, amplitude and duration of the beta-band bursts show progressive changes across trial epochs that explain the temporal pattern of trial-averaged beta-band changes (Rule et al., 2017, 2018; Little et al., 2019), can be correlated with the strength of sensory evidence supporting different action choices (Little et al., 2019), and can discriminate between correct and incorrect action choices (Little et al., 2019). A major conceptual challenge for future research is how to reconcile the highly dynamic, variable nature of real-time beta-band activity with many of its proposed roles, that all implicitly assume an important degree of quasi-static functional stability during each epoch of a trial.

## Conclusions

We report evidence of human premotor cortical beta-band activity that coincided with a sequence of sensorimotor processing stages when participants used a simple color-location conjunction rule to interpret task-relevant information conveyed by arbitrary, non-biological visual cues to select and prepare for wrist movements to report their decision. These effects included a surge in premotor activity at the onset of the task trials, followed by a gradual decrease of premotor activity during the execution of the report movement, with its completion marked by a strong rebound of premotor activity. These modulations over time of premotor activity are compatible with the processing of sensory information to issue a motor plan. We also observed a similar pattern of progressive changes in beta-band activity while the subjects passively observed the same sequence of stimulus events and made a categorical decision about whether a computer applied the color-conjunction rule correctly or not. These responses were seen both in subjects who learned the color-location conjunction rule by trial-and-error while actively performing the task, and in subjects show learned the rule passively while observing a computer perform the task correctly and incorrectly, without associating the visual events with any specific overt wrist movements. This suggests that premotor cortex circuits that normally contribute to the control of voluntary movement based on sensory information from the environment can also contribute to the cognitive interpretation of observed events that should respect specific stimulus-response rules. Finally, differences in beta-band activity between observed correct and incorrect stimulus events further implicate premotor cortex in the assessment of the correctness of observed non-biological visual events.

## Acknowledgements

This work was supported by grants from the Canadian Institutes of Health Research (CIHR MOP-97944 (J.K.) and MOP-142220 (J.K., S.B.), a doctoral studentship from the Natural Sciences and Engineering Research Council of Canada (NSERC)-CREATE: Complex Dynamics of Brain and Behaviour, a scholarship from McGill University Integrated Program in Neuroscience and a doctoral studentship from Fonds de Recherche du Quebec (FRQ). S.B. acknowledges the support from the National Science and Engineering Research Council (NSERC RGPIN 2020-06889), the National Institutes of Health (R01 EB026299-05),the CIHR Canada Research Chair in Neural Dynamics of Brain Systems, the Brain Canada Foundation with support from Health Canada, and the Innovative Ideas program from the Canada First Research Excellence Fund, awarded to McGill University for the Healthy Brains for Healthy Lives initiative.

